# Nucleosome topology and DNA sequence modulate the engagement of pioneer factors SOX2 and OCT4

**DOI:** 10.1101/2022.01.18.476780

**Authors:** Fabiana C. Malaga Gadea, Evgenia N. Nikolova

**Affiliations:** Department of Biophysics, Johns Hopkins University, 3400 N. Charles Street, Baltimore, MD 21218

## Abstract

Nucleosomes in eukaryotic genomes present a barrier to the competent binding of many regulatory proteins. Pioneer transcription factors (pTFs) can bind their target sites on nucleosomal DNA and collaborate with other factors to locally open chromatin and promote transcription. While the interaction of pluripotency pioneer factors and functional partners Sox2 and Oct4 with nucleosomes has been widely studied, molecular details about their engagement in different nucleosome contexts remain elusive. Here, using high-resolution nuclear magnetic resonance (NMR) spectroscopy and biochemical studies, we reveal site-specific structural and dynamic information about pTF interaction with nucleosomes. We find that the affinity of Sox2 and Oct4 to the nucleosome and their synergistic binding correlates with solvent-exposed sites but is highly position and DNA sequence dependent and linked to distinct pTF conformation and dynamics. Sox2 alone forms a super-stable complex near superhelical location 5 (SHL5) with similar affinity and conformation to that of naked DNA but shows elevated dynamics at suboptimal positions. Oct4 strongly favors positions near SHL5.5 and SHL6.5 and both of its DNA binding modules, POU_S_ or POU_HD_, are required for stable complex formation. A ternary complex forms efficiently on canonical Sox2-Oct4 composite motifs (no spacing) near nucleosome ends but is sparse at spaced motifs and absent at internal sites. Moreover, the ability of Sox2 to fold and bend DNA plays a key role in the formation of a stable nucleosome complex and cooperative Oct4 binding. Collectively, our findings describe diverse binding modes of Sox2 and Oct4 on nucleosomes that could guide their site selection and potential interaction with other chromatin factors *in vivo*.

## Introduction

The genomes of eukaryotic organisms are hierarchically organized into a three-dimensional structure that plays critical roles in physiology and disease. The basic structural units of eukaryotic DNA packaging are nucleosomes, which contain ~145 base pairs of DNA wrapped nearly two times around an octamer of four core histone proteins (H2A, H2B, H3, and H4) in a left-handed spiral.^1, 2^ The positioning of nucleosomes on genomic DNA can significantly interfere with the action of proteins called transcription factors (TFs).^3, 4^ TFs recognize specific DNA sequences at the regulatory (promoter and enhancer) regions of genes and ensure correct patterns of gene expression.^5^ Nucleosomes can severely limit access of TFs to their target sites and present a strong barrier to transcription initiation.^6, 7^ To overcome this barrier, cells dynamically modulate nucleosomes through the concerted action of DNA-binding proteins, chromatin modifying and remodeling enzymes, and the RNA Polymerase II transcriptional machinery.^3, 8^

Unlike many TFs, a subset termed “pioneer” TFs (pTFs) can efficiently bind their recognition motifs on DNA embedded in nucleosomes at certain genomic regions.^9, 10^ Pioneer TF binding can lead to opening of silent chromatin regions and trigger diverse transcriptional programs.^11–13^ Genome-wide studies support an intricate mechanism of pTF-induced chromatin opening that involves multiple events and the action of other chromatin factors.^14–16^ Recent high-resolution structural studies show how certain pTFs engage with nucleosomes and possibly perturb the structure of chromatin.^17, 18^ The current consensus model is that pTFs partially unwrap nucleosomal DNA and enable other TFs to bind and unravel nucleosomes, with the help of chromatin remodelers and modifiers, thus promoting transcription.^12, 19^ Still, mechanistic details on how pTFs alter the conformation and packaging of nucleosomes, and if they do so directly or by modulating the activity of proteins that modify and reposition nucleosomes, remain elusive.

Pioneer TFs are represented by FoxA, p53, GATA, and the pluripotency factors Sox2, Oct4, and Klf4, which are known to play critical roles in development, differentiation, and cell reprogramming.^13, 20–23^ These proteins can be amplified, mutated or mis-regulated in cancer and neurological diseases and their aberrant activation or repression can disrupt the architecture of chromatin and rewire gene networks.^24–27^ *In vitro* biochemical surveys as well as *in vivo* genetic studies have uncovered many structurally distinct TFs capable of binding to nucleosomes at different positions and with a range of affinities.^21–23, 28^ TFs with pioneering activity, the so-called nucleosome displacing factors (NDFs), have also been described in budding yeast.^29^ Despite their diverse structures, many pTFs seem to share similar features. They typically recognize DNA motifs via short α-helices in the major groove, which are compatible with nucleosome topology and, thus, are not be expected to significantly alter nucleosome structure.^23^ A well-studied representative of this group is human FoxA, which structurally resembles the linker histone H1.^30^ FoxA binds preferentially to its motif near the nucleosome dyad^11^ and maintains local chromatin accessible by competitively displacing H1.^31^ Other modes of DNA recognition by pTFs include HMG box domains, which bind in the DNA minor groove and induce a large helical bending.^32^ HMG proteins, including Sox2, are believed to favor nucleosome binding by exploiting the pre-bent conformation of DNA.^23, 33^

How do pTFs recognize nucleosomes and target select sites in the genome? In the nucleosome, DNA curvature and histone contacts modify the physico-chemical properties and accessibility of the DNA helix. DNA bending leads to widening of the major and minor groove facing outward and narrowing of the grooves facing in towards the histone core.^1^ Multiple studies suggest that pTFs prefer the solvent-exposed, outside surface and disfavor the buried, inner surface of nucleosomal DNA.^17, 18, 28, 34^ Direct structural insights into pTF-nucleosome recognition come from a limited number of high-resolution studies of Sox2 alone or with its functional partner Oct4 obtained by cryo-electron microscopy (cryo-EM).^17, 18^ Sox2 and Oct4 are pTFs that regulate cell-fate decisions^35^ and are known to engage with silent chromatin in embryonic stem cells (ESCs) and during induced pluripotent stem cell (iPSC) generation.^33, 36, 37^ They can bind synergistically and interact directly on adjacent sites *in vitro*^38^ and colocalize on target genes *in vivo*.^39–41^ To perform their synergistic role, Sox2 and Oct4 co-bind on adjacent (canonical) or closely spaced motifs, where the spacing determines their interactions and degree of cooperativity.^42, 43^ The Sox2 HMG domain folds into an L-shaped structure and binds DNA in the minor groove, severely bending and unwinding the helix.^42, 44^ Oct4 recognizes a longer octameric motif using a bi-partite domain comprised of a POU-specific (POU_S_) and a POU-homeodomain (POU_HD_) subunit connected by a flexible linker.^42, 44, 45^ On the canonical composite motif, found in the Nanog promoter, protein-protein contacts and shape complementarity occur between the Sox2 HMG α3 helix and the POU_S_ subunit.^43^ On the 3 base-pair spaced composite motif, represented by the Fibroblast growth factor 4 (Fgf4) enhancer, contacts are formed between the HMG C-terminal tail and POU_S_ and confer weaker cooperativity.^42^ The HMG domain ends up on the same face of the DNA helix as POU_S_ on Nanog (canonical) sites and as POU_HD_ on Fgf4 sites. This configuration predisposes the canonical motif, where both HMG and POU_S_ can bind accessible surfaces, for optimal synergism on nucleosomes.

The cryo-EM studies of Sox2 bound to nucleosomes, with or without Oct4 on a canonical motif, show that the two pTFs bind at adjacent solvent-exposed sites and perturb nucleosome structure to a variable extent.^17, 18^ Binding of Sox2 at an internal site near superhelical location 2 (SHL2) locally deforms DNA and can cause unwrapping of the adjacent gyre.^18^ In the presence of Oct4 at a less internal site (~SHL5), Sox2 induces large DNA bending similar to its naked DNA-bound state without unwrapping from the histone core.^17^ Binding of Sox2 and Oct4 near the DNA entry and exit points (~SHL6) severely distorts the DNA helix, which could displace H1 and affect inter-nucleosome packing.^17^ In the tertiary complexes, Oct4 engages nucleosomal DNA with only one of its DNA-binding modules (POU_S_) at its solvent-exposed half-site, while the half-site for POU_HD_ remains occluded by the histone core.^17^ The POU_HD_ subunit is not observed and, as suggested by molecular dynamics (MD) simulations,^46, 47^ may adopt other configurations and transiently interact with both DNA and histones. Although these studies have provided crucial insights in pTF-nucleosome recognition, a systematic analysis of the binding affinity, protein conformation, and synergism of Sox2 and Oct4 as a function of nucleosome position, composite motif and nucleosomal DNA sequence is needed. Also, the dynamic nature of Sox2 and Oct4 binding modes and concurrent nucleosome perturbations cannot be adequately visualized using X-ray crystallography and cryo-EM techniques. We propose to address these challenges by using state-of-the-art solution NMR spectroscopy methods tailored for large biomolecules (methyl TROSY)^48, 49^ to interrogate pTF conformation and dynamics as well as pTF-induced structural and dynamics changes in nucleosomes. Here, we combine traditional biochemistry, chemical probing and powerful NMR techniques to determine the binding affinities and the conformation of Sox2 and Oct4 for a range of nucleosome positions and in the context of different composite motifs. Our findings suggest that the affinity and synergistic binding of Sox2 and Oct4 to nucleosomes correlate with solvent-exposed sites but are highly position and DNA sequence dependent and can be linked to different pTF conformation and dynamics. These binding preferences are likely guided by variations in histone-DNA contacts, DNA shape and flexibility across the nucleosome.

## Results

### Design of nucleosome constructs with Sox2 and Oct4 composite binding sites

In this study, we aimed to determine the binding affinity of Sox2 and Oct4 to nucleosome core particles (NCPs) and its modulation by nucleosome position and composite motif sequence. Towards this, we prepared double fluorophore-labeled (5’-FAM and 5’-Cy3) mono-nucleosomes with the Sox2-Oct4 composite site located at variable positions along a 145 base-pair fragment with the Widom 601 nucleosome positioning sequence (Figure 1A, Table S1).^50^ The original 601 sequence was modified on one end (601*) to replace a degenerate Sox2-binding site in nucleosomes (see Table S1). The Sox2-Oct4 composite site was derived from the regulatory enhancer region of the mouse Fgf4 (mFgf4)^51^ and the proximal promoter region of the mouse Nanog (mNanog)^39^ genes involved in embryonic development (Figure 1A, B). The mFgf4 motif contains a 3 base-pair spacer between the Sox2 (italicized) and Oct4 (underlined) binding sites (5’*CTTTGTT*TGG<u>ATGCTAAT</u>), while the mNanog motif contains no spacing between the two binding sites (5’*CATTGTA*<u>ATGCAAAA</u>) (Figure 1A, B). The composite motifs were typically placed in a way to ensure the DNA minor groove of the Sox2 site was facing outward and away from the histone core and the second DNA gyre, oriented in the 5’-to-3’ direction on the FAM-labeled strand (NCP 2R, 23R, 54R, 55R, and 65R) or the Cy3-labeled strand (NCP 0, 52, and 62) (Figure 1A). The naming of the NCP (DNA) constructs reflects the location of the central G-C base-pair within the Sox2 motif (5’*CT/ATT<u>G</u>TT/A*) on the 601* sequence and the subscript identifies the mFgf4 (F) or mNanog (N) motif. The motifs were inserted in the forward or reverse orientation at the dyad and near SHL2, 3, 5 and 6 (see Figure 1A). Several other sites were designed so that the Sox2 motif was only partially exposed (NCP 27, 32). The 601* sequence was used as a control. The accessibility of the Oct4 major groove binding site varied depending on the inter-motif spacing and rotational setting on the nucleosome (Figure 1A). Specifically, the POUs subunit binding site (5’ ATGC) was mostly accessible in 2R_N_, 23R_N_, 32_F_, 54R_N_, and 62_N_, while the POU_HD_ subunit binding site (5’ T/AAAT/A) was mostly accessible in 0_F_, 2R_F_, 23R_F_, 27_N_, 52_F_, 54R_F_, 55R_F_, and 62_F_ constructs. The location of the composite motif in NCP 54R_N_ and 62_N_ constructs was the same (62_N_ was symmetrically flipped to the + side) as in the cryo-EM structures of Sox2 and Oct4 bound to a 601-based nucleosome.^17^

**Figure 1.**
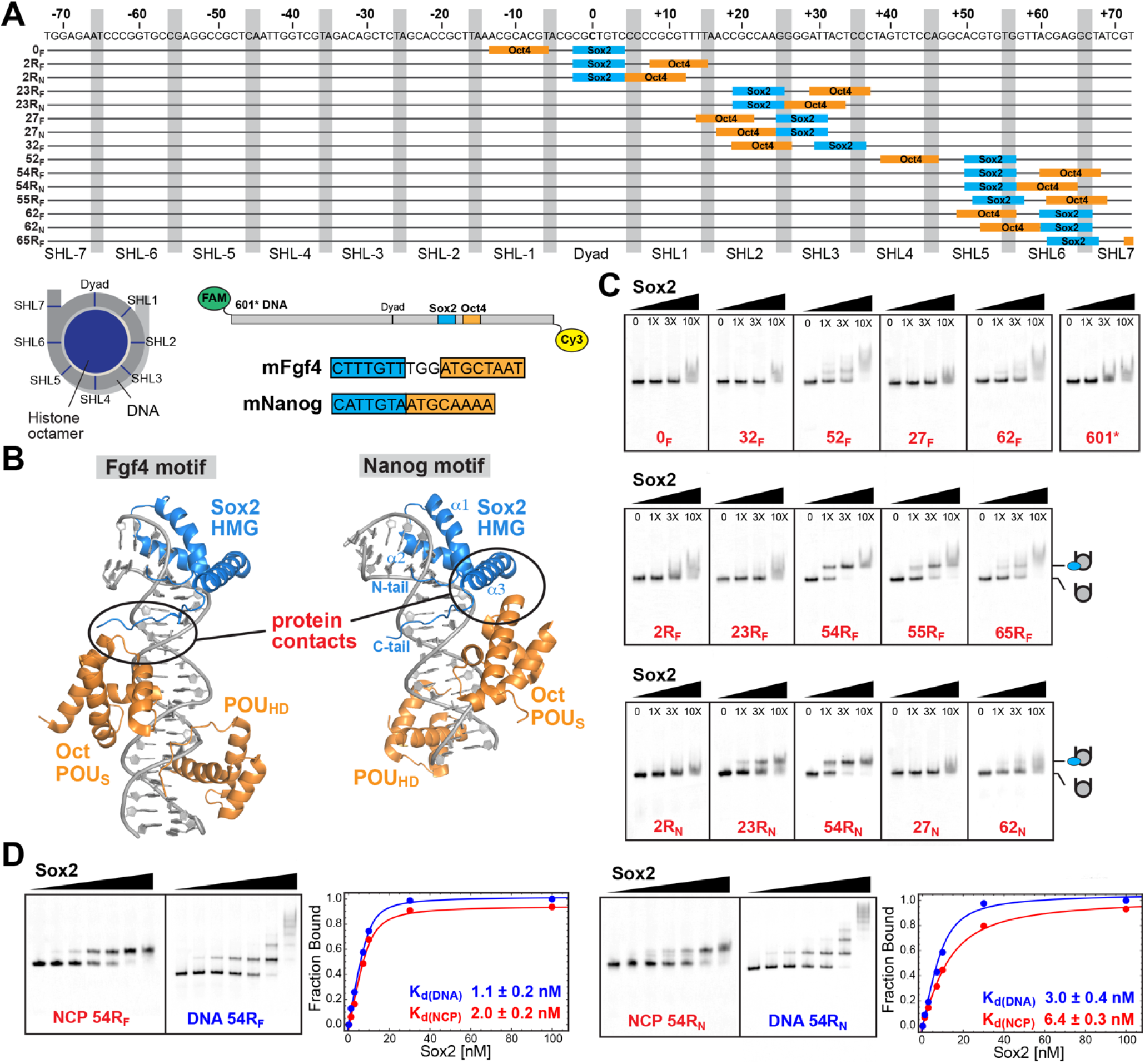
Sox2 binding to nucleosomes favors SHL5 but is highly position and DNA sequence dependent. (A) DNA constructs containing the Sox2-Oct4 composite motif (mFgf4 or mNanog) at different translational and rotational positions on the nucleosome. The motifs were inserted in a 145 base-pair modified Widom 601 sequence (601*, shown on top) that was double-labeled with a 5’-FAM (strand shown) and 5’-Cy3 fluorescence probes for biochemical assays. The nucleotides are numbered from −72 to + 72 around the dyad (0) and each superhelical location (SHL) is indicated below the sequence and on the NCP cartoon. The grey rectangles highlight the narrow minor groove facing towards the histone core, including the strong positioning TpA dinucleotide steps. (B) Models of Sox2 and Oct4 binding to mFgf4 and mNanog composite motifs based on crystal (left, PDB ID: 1GT0) and NMR (right, PDB ID: 1O4X) structures of Sox2-Oct1-DNA (Oct1 is a close homolog of Oct4). (C) EMSA data for the Sox2 HMG domain binding (0-100nM) to different NCP constructs (10nM) showing variations with position and binding motif sequence. The highest affinity complex is formed with NCP 54R for both mFgf4 and mNanog motifs. (C) Representative EMSA titrations of Sox2 HMG (0-100nM) with NCP or DNA 54R constructs (10nM) and corresponding best fits to a two-state binding model with apparent dissociation constants (K_d_, inset). K_d_ values and errors were calculated as the average and standard deviation of at least three independent measurements.

### Sox2 binds distinct nucleosome positions and forms a super-stable complex at SHL5

First, we probed the relative binding affinity of the human Sox2 HMG DNA-binding domain (DBD, residues 39-118) to nucleosomes or naked DNA using electrophoretic mobility shift assay (EMSA) (Figure 1 and Figure S1) and quantified the apparent dissociation constant (K_d_) at near-physiological salt concentrations (Table 1). We observed formation of a discrete super-shifted band representing the Sox2-nucleosome complex at positions 23R, 54R, 55R, and 65R and a less shifted weaker band at positions 52 and 62 (Figure 1C). Sox2 bound most tightly to NCP 54R at a distinct position near SHL5 with a slightly higher affinity for the mFgf4 (54R_F_, K_d_ ~ 2.0 ± 0.2 nM) than the mNanog sequence (54R_N_, K_d_ ~ 6.4 ± 0.3 nM) (Figure 1D). In comparison, Sox2 binding to DNA 54R was only ~2-fold tighter than binding to NCP 54R, with K_d_ of 1.1 ± 0.2 nM and 3.0 ± 0.4 nM for the mFgf4 and mNanog motifs in naked DNA, respectively (Figure 1D). Shifting the composite site by one base-pair in NCP 55R_F_ led to a detectable 4-fold decrease in binding affinity (K_d_ ~ 8.3 ± 1.0 nM) and lower complex stability relative to NCP 54R_F_, as indicated by more pronounced smearing between the free and NCP-bound state (Figure 1C). Translating the composite site to a more internal position near SHL2 in NCP 23R attenuated Sox2 binding at the mNanog motif (K_d_ ~ 12.7 ± 1.3 nM) and completely abolished the discrete complex at the mFgf4 motif (Table 1). Surprisingly, moving the composite motif by one helical turn and closer to the nucleosome edge (entry/exit DNA) in NCP 65R_F_, where transient DNA unwrapping occurs,^52^ diminished the affinity and stability of the complex (K_d_ ~ 12.4 ± 1.2 nM). Alternatively, inverting the orientation of the Sox2 motif while preserving its position on DNA in NCP 52_F_ as compared to NCP 54R_F_ also led to a sizeable decrease in binding (K_d_ ~ 11.1 ± 1.4 nM) and a less shifted band for the complex, possibly signifying a lower degree of DNA deformation. The binding of Sox2 to NCP 62 (K_d_ ~ 45 ± 7 nM for mFgf4 and K_d_ ~ 38 ± 5 nM mNanog) was similarly weakened relative to the inverted site in NCP 65R_F_. Other internal nucleosome positions displayed mostly non-specific binding that was indistinguishable from the control 601* nucleosome (K_d_ ~ 41 ± 4 nM) (Figure 1C, Table 1). Constructs with the Sox2 motifs placed in either orientation at the dyad, where it is largely accessible, showed only marginally stronger binding affinities (i.e. K_d_ ~ 29 ± 3 nM for NCP 0_F_) than the control sequence and substantially weaker affinity than for NCP 54R constructs. The affinities for weak or non-specific binding sites agreed well with previously published fluorescence polarization measurements of Sox2 binding to a similar 601-based nucleosome (K_d_ ~20-60 nM).^17^ At the same time, nanomolar affinities (K_d_ ~0.3-1.4 nM) comparable to those of NCP 54R_F_ have been previously reported for the full-length human Sox2 protein binding to a *Lin28B*-based nucleosome under low salt conditions.^33^ Moreover, while the binding affinities of Sox2 to nucleosomes recorded here were highly sensitive to the superhelical position (K_d_ ~2-50 nM) for solvent-exposed motifs, the affinities for the corresponding naked DNA constructs were much less variable (K_d_ ~1-4 nM). Our findings point to a strong position and sequence preference of Sox2 association with nucleosomes, which is likely guided by the nucleosome shape and histone-DNA contacts.

**Table 1.** Apparent binding affinities (K_d_) of Sox2 or Oct4 for different DNA and NCP constructs containing the mFgf4 or mNanog site.

| Construct | $K_d$ (nM) | |
| --- | --- | --- |
|  | Sox2 | Oct4 |
| ***mFgf4*** |  |  |
| DNA 2R<sub>F</sub> | 3.3 ± 0.5 |  |
| DNA 54R<sub>F</sub> | 1.1 ± 0.2 | 32 ± 3 |
| DNA 62<sub>F</sub> | 2.1 ± 0.2 |  |
| NCP 2R<sub>F</sub> | 31 ± 4 |  |
| NCP 23R<sub>F</sub> | 37 ± 4 |  |
| NCP 54R<sub>F</sub> | 2.0 ± 0.2 | 240 ± 30 |
| NCP 55R<sub>F</sub> | 8.3 ± 1.0 | 110 ± 10 |
| NCP 65R<sub>F</sub> | 12.4 ± 1.2 |  |
| NCP 0<sub>F</sub> | 29 ± 3 |  |
| NCP 27<sub>F</sub> | 34 ± 4 |  |
| NCP 32<sub>F</sub> | 24 ± 3 |  |
| NCP 52<sub>F</sub> | 11.1 ± 1.4 |  |
| NCP 62<sub>F</sub> | 41 ± 7 | 390 ± 50 |
| ***mNanog*** |  |  |
| DNA 2R<sub>N</sub> | 3.7 ± 0.5 |  |
| DNA 54R<sub>N</sub> | 3.0 ± 0.4 | 38 ± 4 |
| DNA 62<sub>N</sub> | 2.3 ± 0.3 |  |
| NCP 2R<sub>N</sub> | 23 ± 3 |  |
| NCP 23R<sub>N</sub> | 12.7 ± 1.3 |  |
| NCP 54R<sub>N</sub> | 6.4 ± 0.3 | 250 ± 20 |
| NCP 27<sub>N</sub> | 33 ± 4 |  |
| NCP 62<sub>N</sub> | 38 ± 5 | 110 ± 10 |
| ***No motif*** |  |  |
| DNA 601* | 8.0 ± 0.7 | 160 ± 20 |
| NCP 601* | 41 ± 4 | 350 ± 40 |

### Oct4 forms a stable lower-affinity nucleosome complex near SHL5.5 and SHL6.5

Next, we used EMSA to interrogate the binding of the Oct4 bipartite DBD, which consists of a POU_S_ and POU_HD_ subunit connected by a long flexible linker, to DNA and nucleosomes (Table 1). In agreement with prior studies,^17, 46, 53^ Oct4 alone formed a discrete stable complex only with nucleosomes that contained the binding site near SHL5.5 and SHL6.5, including NCP 54R_F_, 55R_F_, 54R_N_ and 62_N_ (Figure 2 and Figure S1). Two super-shifted bands were observed with NCP 55R_F_, 54R_N_, and 62_N_ at low ratios of Oct4 to NCP (Figure 2A and Figure S1). This suggests that Oct4 may interact with more than one site on these constructs or utilize distinct modes of binding to its cognate motif that yield different gel migration. In addition, we note that Oct4 can bind stably to nucleosomes, where the POU_S_ motif is exposed and POU_HD_ motif occluded (NCP 54R_N_ and 62_N_) and vice versa (NCP 54R_F_ and 55R_F_) (Figure 2A, C). This behavior could stem from the potential of either Oct4 subunit to associate independently with DNA (Figure S2). We also observed a small amount of similarly super-shifted bands for NCP 52_F_ and 62_F_, which disappeared at high Oct4 ratios, likely due to low complex stability and non-specific competition (Figure 2B). Quantitative analysis of the EMSA data revealed that the high-affinity binding mode of Oct4 to nucleosomes was substantially weaker than that of Sox2 (Table 1). The association of Oct4 was slightly stronger with NCP 62_N_ (K_d_ ~ 110 ± 10 nM) and 55R_F_ (K_d_ ~ 110 ± 10 nM) as compared to NCP 54R_F_ (K_d_ ~ 240 ± 30 nM) and 54R_N_ (K_d_ ~ 250 ± 20 nM). By contrast, association of Sox2 with NCP 54R_F_ and 54R_N_ was roughly 100-fold and 40-fold tighter, respectively. Moreover, Oct4 binding to DNA comprising the mFgf4 (K_d_ 32 ± 3 nM) or mNanog (K_d_ 38 ± 4 nM) motifs was significantly tighter than binding to the respective NCPs. All other nucleosomes formed a relatively unstable complex with Oct4 with a substantially lower affinity comparable to the control sequence (Figure 2 and Figure S1).

**Figure 2.**
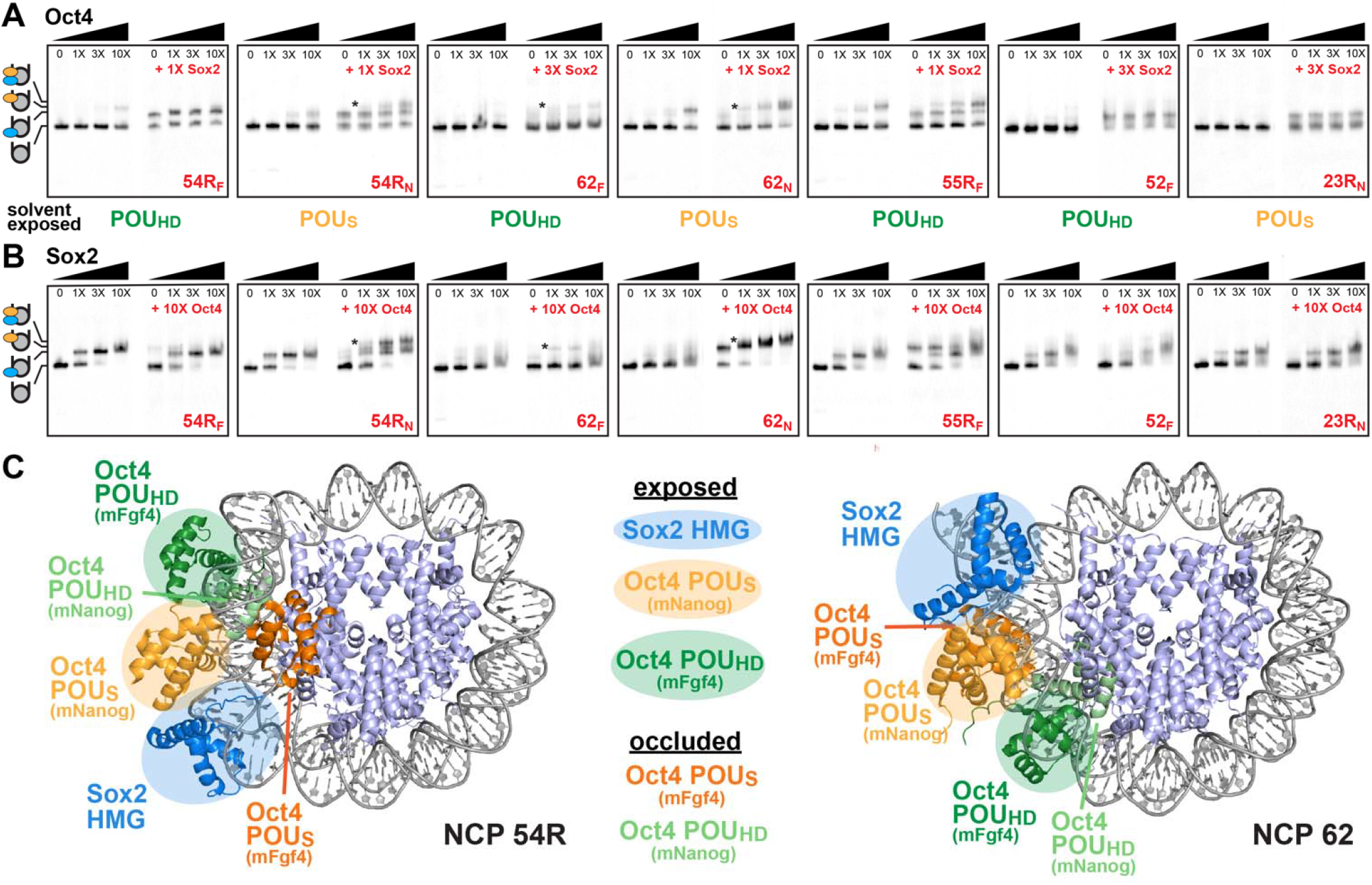
Nucleosome position and spacing of the composite motif affect Oct4 binding and Sox2-Oct4 synergism. (A) EMSA of Oct4 (0-100nM) binding alone or in the presence of Sox2 (10 or 30nM) to NCP constructs (10nM). The Oct4 subunit, POU_S_ or POU_HD_, whose half-site is solvent exposed in each NCP construct is highlighted below each gel. (B) EMSA of Sox2 (0-100nM) binding alone or in the presence of Oct4 (100nM) to NCP constructs (10nM). The tertiary Sox2-Oct4-NCP complex is observed to variable extent for certain composite motifs near SHL5 and 6, denoted by an asterix (*), and not observed for more internal sites. (C) Overlay of the cryo-EM structures of Sox2-Oct4 bound to a nucleosome at the same positions as in NCP 54R (PDB ID: 6YOV) and in NCP 62 (PDB ID: 6T90) with the NMR or crystal structure of the Sox2-Oct1-DNA complex on a canonical (PDB ID: 1O4X) or the mFgf4 (PDB ID: 1GT0) composite motif. For mNanog (canonical) motifs, Sox2 HMG (blue) and Oct4 POU_S_ (light orange) are predicted to bind the outer-facing DNA surface and can synergize, while Oct4 POU_HD_ (light green) is predicted to bind the occluded DNA surface and likely adopts an alternative conformation. For mFgf4 motifs, Sox2 HMG and Oct4 POU_HD_ (dark green) are predicted to bind the exposed DNA surface and Oct4 POU_S_ (dark orange) is predicted to overlap with the histone core (NCP 54R) or other DNA gyre (NCP 62). There, cooperative binding would require partial DNA dissociation.

To dissect the ability of each Oct4 module to interact independently with nucleosomes, we also probed the binding of truncated protein constructs corresponding to the POU_S_ or POU_HD_ subunits to nucleosomes (Figure S2). Unlike with the full Oct4 DBD, binding of POU_S_ or POU_HD_ to NCP 54R_F_ and 54R_N_ did not result in a distinct band but instead a super-shifted smear. Also, POU_S_ appeared to be saturating NCP 54R_F_ at lower concentrations than POU_HD_. This was unexpected, as the POU_S_ half-site in mFgf4 is predicted to be obstructed by histones, while the POU_HD_ half-site should remain accessible (Figure 2C). This signifies that POU_S_ might be interacting with a different sequence within the mFgf4 motif in NCP 54R_F_ (and 55R_F_). Lastly, simultaneous addition of POU_S_ and POU_HD_ to either NCP 54R_F_ or 54R_N_ did not recapitulate the binding profile of the full Oct4 DBD. In fact, it appeared to inhibit binding relative to each truncated protein alone, which could imply possible competition or aggregation. Thus, our results show that the linked configuration of the POU_S_ and POU_HD_ sub-domains within the Oct4 DBD is required for high-affinity binding at SHL5.5 and SHL6.5. These findings are in line with recent reports showing diminished Oct4-nucleosome binding with alterations in the linker region.^54^

### Sox2-Oct4 synergism is enhanced on canonical motifs near nucleosome edges

We further employed EMSA to monitor how nucleosome position affects the formation of the Sox2-Oct4-NCP ternary complex and the synergistic action of Sox2 and Oct4 (Figure 2A, B and Figure S3). Most prominently, we observed a clear stabilization of Sox2 or Oct4 binding to the nucleosome in the presence of the other protein for NCP 54R_N_ and, more strongly, to NCP 62_N_ containing the canonical mNanog motif (Figure 2A, B). The ternary complex appeared as a super-shifted band migrating slower than the binary complexes in NCP 54R_N_ or as an overall more retarded band in NCP 62_N_. The different migration pattern for NCP 62_N_ could be due to faster binding kinetics and/or different conformations with similar electrophoretic mobilities. Stable formation of the ternary complex was not surprising in mNanog constructs, where the orientation of the Sox2 HMG and Oct4 POU_S_ sites made them accessible and enabled direct interaction between the two proteins. However, the ternary complex could not be detected on a more internal position in NCP 23R_N_, where both the Sox2 and Oct4 POU_S_ sites were still solvent exposed. This suggests that factors other than solvent accessibility of nucleosomal DNA influence the cooperative binding of the two proteins.

By contrast, the ternary complex was scarce or entirely absent on nucleosomes containing the spaced mFgf4 motif, consistent with the POU_S_ half-site being occluded by the histone octamer (Figure 2C). For NCP 54R_F_ and 55R_F_, we detected two resolved bands in the presence of both proteins, which corresponded to the binary Sox2-NCP and Oct4-NCP species (Figure 2A, B). The Oct4-NCP signal disappeared as the concentration of Sox2 was increased, possibly with non-specific binding or allosteric interactions displacing Oct4 from the nucleosome. Interestingly, for NCP 62_F_ that does not associate stably with either Sox2 or Oct4, we observed a small but detectable amount of a new super-shifted band consistent with a ternary Sox2-Oct4-NCP complex. This band was not detectable in NCP 52_F_, where the mFgf4 motif is placed in a similar rotational setting as in NCP 62_F_ but translated internally by one helical turn. Finally, the ternary complex was not observed near the dyad (NCP 0, 2R) or at an internal site (NCP 27) containing either the mFgf4 or mNanog motif or in the control sequence (Figure S3). These results suggest that the ternary complex is disfavored at internal positions, regardless of the motif sequence or spacing between the Sox2 and Oct4 sites. However, it can be efficiently formed at canonical composite sites (mNanog) near SHL5 and SHL6 or transiently established at spaced composite sites (mFgf4) incorporated at nucleosome ends, possibly due to spontaneous DNA unwrapping.

### Sox2 binding at SHL5 mimics DNA-bound state but is dynamic at suboptimal positions

Next, to gain insight into the conformation of various Sox2-NCP complexes, including the super-stable complex at SHL5, we employed chemical and enzymatic probing coupled with NMR experiments tailored for super-large biomolecules. First, we used DNaseI footprinting of Sox2 and Oct4, alone or together, with various NCPs and corresponding naked DNA to pinpoint the location of the bound protein, which would manifest as reduced nuclease cleavage (Figure 3 and Figure S4). First, we observed a strong footprint of Sox2 at its cognate motif on naked DNA (any construct) as well as weaker footprints at other non-cognate sites, consistent with the non-specific binding seen by EMSA (Figure 1 and Figure S1). For nucleosomes, Sox2 binding to NCP 54R_F_ and 55R_F_ also resulted in a selective DNaseI footprint at the mFgf4 motif, while no reduced cleavage was detected on other sites (Figure 3A). A more pronounced footprint was observed for NCP 54R_F_ than 55R_F_, consistent with the higher affinity of Sox2 determined here. Oct4 binding to naked DNA generated a single footprint near its cognate motif, which was slightly enhanced with the addition of Sox2 (Figure 3A and Figure S4). A weaker reduction of DNaseI cleavage near the Oct4 motif was also observed in the context of nucleosomes. A stronger footprint for NCP 55R_F_ again correlated with tighter Oct4 binding at that site. Weaker Oct4 binding as compared to Sox2 and high nuclease activity near the Oct4 motif at the DNA termini most likely contributed to the weaker Oct4 footprint relative to that of Sox2. By contrast, the footprint could be clearly detected for internal Oct4 motif positions in naked DNA (Figure S4). These results indicate that both Sox2 and Oct4 recognized their specific motifs on naked DNA and can bind their motifs selectively in the context of nucleosomes.

**Figure 3.**
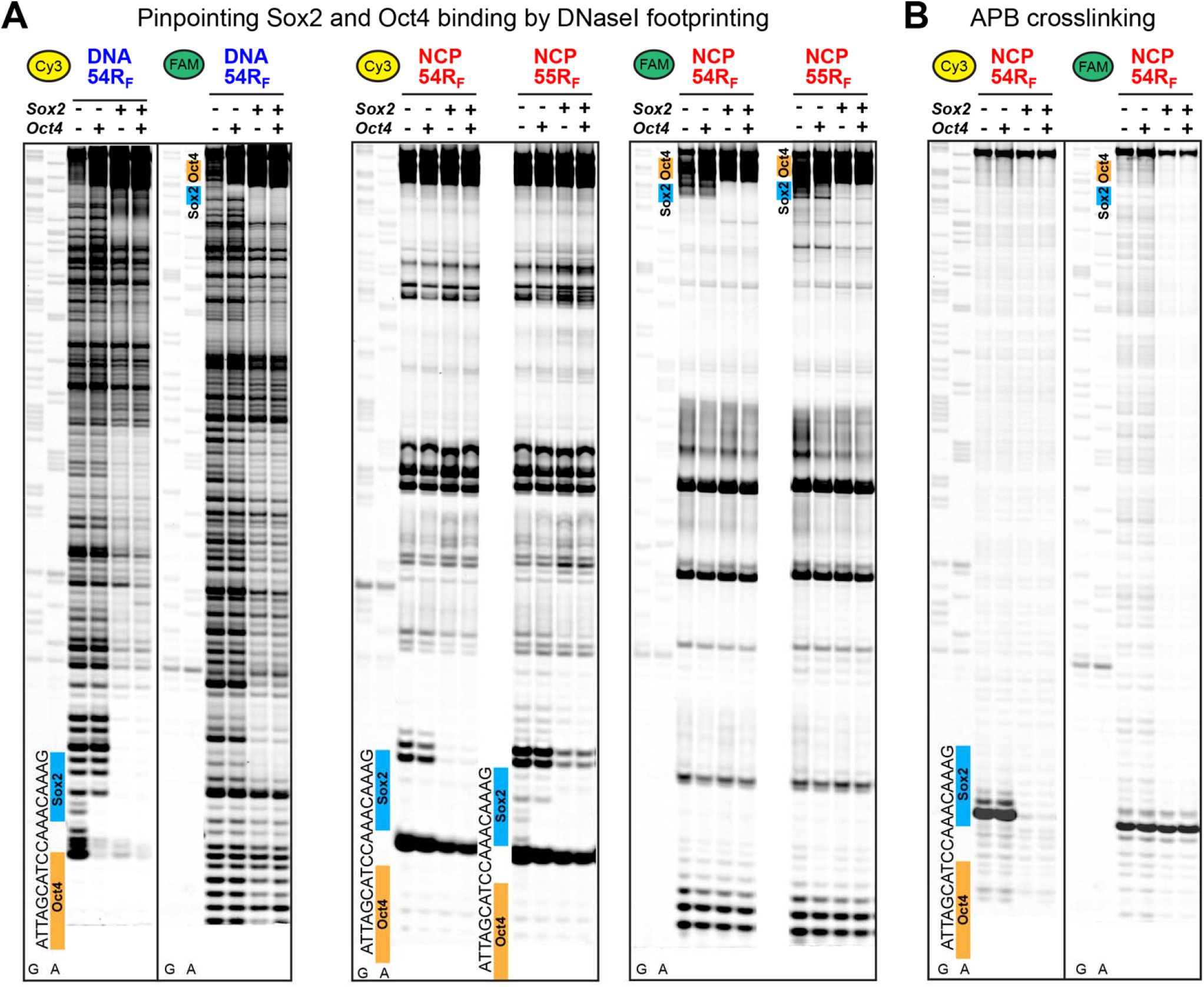
The high-affinity Sox2-nucleosome complex at SHL5 features a large DNA deformation. (A) DNaseI footprinting assay of Sox2 and Oct4 (1μM each) binding to DNA 54R_F_ (100nM) or NCP 54R_F_ and 55R_F_ (100nM). The position and sequence of the Sox2 (blue) and Oct4 (orange) site on the mFgf4 composite motif is indicated on each gel. A clear footprint for Sox2 and Oct4 near their motifs on free DNA indicates site-specific binding (Sox2 shows non-specific binding at other sites). A clear footprint for Sox2 near its motif in NCP shows stable complex formation at SHL5. The Oct4 footprint on NCP is not clearly observed due to proximity to DNA ends and/or faster kinetics. (B) APB crosslinking assay of Sox2 and Oct4 (1μM each) binding to NCP 54R_F_ H2B(S53C) (100nM) showing suppression of the histone-DNA crosslink in the presence of Sox2 (but not Oct4). This is consistent with a large protein-induced DNA deformation. (DNA ladders for G and A nucleotides may vary slightly in sequence from constructs shown in gels.)

Next, we used chemical crosslinking to probe whether Sox2 binding alone induced a large deformation in nucleosomal DNA similar to those observed in its complex with naked DNA and in the ternary complex with Oct4 on nucleosomes. Towards this, we prepared an NCP 54R_F_ construct with histone H2B harboring a cysteine mutation at Serine 53 (S53C) in a cysteine-free histone background (H3(C110A)). The sulfhydryl group on the cysteine residue was chemically labeled with the photolabile crosslinker 4-azidophenacyl bromide (APB). When exposed to UV light, the APB moiety crosslinks with nucleosomal DNA in its proximity (~10 Å).^55, 56^ The H2B(S53C), when functionalized with APB on both H2B histones, has been shown to effectively crosslink to DNA (positions ±53 and ±55, Figure 1A) on both sides of the nucleosome.^56^ Based on the structure of the Sox2-Oct4-NCP ternary complex, where a large DNA helical bending pulls the backbone away from H2B(S53) by over 8 Å, we predicted that APB crosslinking would be suppressed upon Sox2 binding. Here, we carried out a crosslinking reaction of NCP54_F_ H2B(S53C) in the presence or absence of Sox2 and Oct4. Nucleosomal DNA, selectively cleaved at the crosslinked sites, was visualized on both strands on a sequencing gel at base-pair resolution. In line with previous studies, we observed pronounced cleavage in free nucleosomes at position +55 and −55 on the on the Cy3-labeled and FAM-labeled DNA strand, respectively (Figure 3B). Addition of Sox2 abolished DNA cleavage at base-pair +55 (Cy3 strand) located within the mFgf4 motif but not at base-pair −55 (FAM strand) that lacks the Sox2 motif. DNA cleavage was reduced to a similar degree with Sox2 alone or in the presence of Oct4. By contrast, addition of Oct4 only to NCP54_F_ H2B(S53C) did not decrease the extent of DNA cleavage as compared to the free NCP. Our data strongly suggests that Sox2 binding alone induces severe deformation of nucleosomal DNA, likely similar to that observed in Sox2-DNA and Sox2-Oct4-NCP complexes. Oct4, on the other hand, does not appear to considerably distort nucleosomal DNA near the crosslink site but may still cause DNA deformation several base-pairs away.

We also wondered if the conformation of Sox2 bound to NCP54_F_ was similar to that of the DNA-bound state. To this end, we used solution NMR spectroscopy methods tailored for large biomolecules to probe in atomic detail the conformation of Sox2 in complex with NCP54_F_ (molecular weight (MW) ~200 kDa) and other lower-affinity NCP constructs. The observation of high-MW complexes by solution-state NMR using conventional uniform isotopic labelling is typically hindered by the slow overall tumbling and fast transverse relaxation rates, leading to extreme line broadening of the resonance signals. Methyl transverse relaxation-optimized spectroscopy (TROSY)-based methods take advantage of the slowly relaxing coherences of methyl (^13^CH_3_) groups in large biomolecules to detect and characterize super-large protein and nucleic acid complexes with improved resolution and sensitivity.^48, 49^ Here, we employed site-specific ^13^C-labeling of isoleucine, leucine, valine, and alanine (ILVA) methyl groups of the Sox2 HMG DNA-binding domain in a deuterated protein background combined with methyl TROSY NMR to probe Sox2 binding to nucleosomes or naked DNA.^57^ Sox2 HMG methyl resonance assignments for the free and DNA-bound state were obtained using a ^13^C, ^15^N-labeled protein alone or in complex with a short (15 base-pair) DNA duplex containing the mFgf4 site (sDNA_F_). Figure 4A shows a methyl ^1^H,^13^C-HMQC spectral overlay of Sox2 free, bound to sDNA_F_ or bound to NCP 54R_F_. Sox2 association with its cognate DNA motif causes sizeable chemical shift changes in a number of methyl sites near the protein-DNA interface (Figure 4A), which can be used as reporters of the Sox2 conformation. We also confirmed that the complexes with short and long (145 base-pair) DNA are nearly identical (Figure S5). For comparison, Sox2 binding to a random non-specific short DNA (sDNA_Ctr_) leads to distinct chemical shift changes and significant line broadening for the respective sites (Figure S5). When examining nucleosome complexes, we found that the Sox2-NCP 54R_F_ complex resembles closely the Sox2-DNA bound state, with only small chemical shift changes observed for the NCP-bound state. Several additional weaker peaks were visible in Sox2-NCP 54R_F_ that overlay with peaks of the free Sox2 (i.e. V5, A11, V14, L45), which could be due to incomplete saturation or alternative bound conformations of Sox2. Our findings that the Sox2 conformation is similar for specific DNA and NCP 54R_F_ complexes is consistent with the similar high affinities for these constructs obtained by EMSA and the nucleosomal DNA deformation observed in crosslinking experiments.

**Figure 4.**
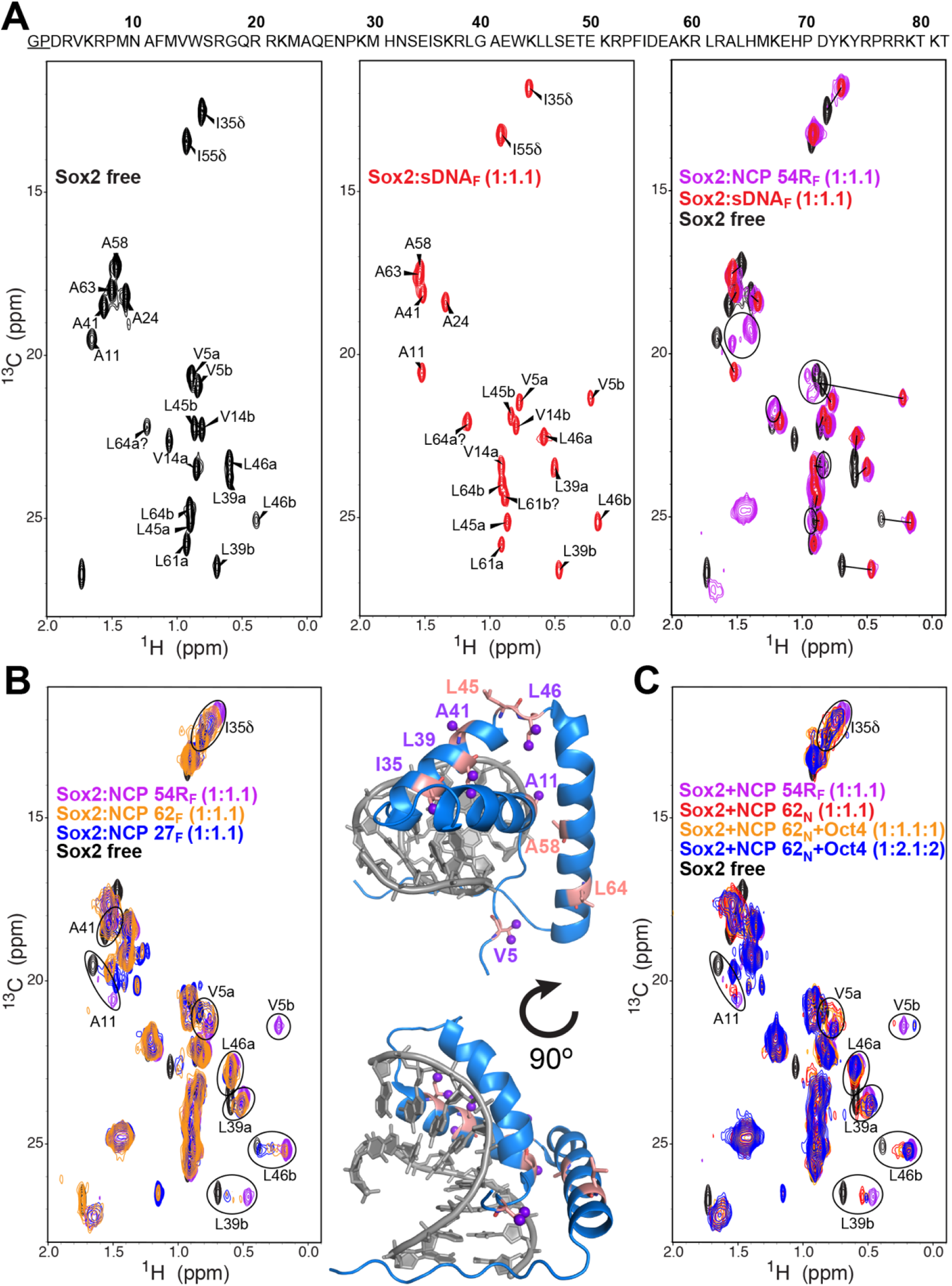
The conformation of Sox2 bound at SHL5 mimics the stable DNA-bound state but is dynamic at suboptimal nucleosome positions. (A) ^1^H, ^13^C-HMQC methyl NMR spectra of ILVA-labeled Sox2 HMG domain (residues 3-82) in the free state (black) and bound to a short DNA (red, sDNA_F_) or NCP 54R_F_ (violet) containing the mFgf4 motif. Assignments for the Sox2 free and DNA-bound state are indicated. The spectral overlay (right) shows similar chemical shifts between the DNA- and NCP-bound states of Sox2 and large chemical shift changes relative to the Sox2 free state. Some peaks in the Sox2-NCP 54R_F_ complex (circled) match peaks for the Sox2 free state and could reflect incomplete saturation or alternative conformations. (B) Overlay of ^1^H, ^13^C-HMQC spectra of the ILVA-labeled Sox2 bound to nucleosomes with different binding affinities (left) showing that certain residues (circled) in NCP 27_F_ and 62_F_ complexes occupy chemical shifts between the Sox2 free and DNA-bound state. Structure of Sox2 bound to DNA (PDB ID: 1GT0) highlighting residues that undergo large chemical shift changes between the free and DNA-bound state (salmon) shown in (A). Methyl groups that exhibit chemical shift changes between different NCP complexes in spectra on the left (circled) are displayed as purple spheres and coincide with residues in salmon. (C) Overlay of ^1^H, ^13^C-HMQC spectra of the ILVA-labeled Sox2 bound to NCP 62_N_ in the presence and absence of Oct4. The tertiary Sox2-Oct4-NCP complex near SHL6 shows persistent line broadening and chemical shift differences relative to the high-affinity Sox2-NCP 54R_F_ complex (violet) with excess Oct4 and NCP.

We further investigated the complex between Sox2 and weaker binding nucleosomes, in the presence or absence of Oct4. For NCP 62_F_ and 62_N_, a number of residues in the 1:1 complex with Sox2 displayed significant line broadening and multiple corresponding peaks residing between the chemical shift of the Sox2 free and DNA-bound state (Figure 4B). For NCP 27_F_, which binds even less tightly to Sox2, we observed a similar behavior. However, the peaks were shifted closer to the protein free state in the 1:1 complex and moved only slightly towards to the DNA-bound state with excess of nucleosome (Figure 4B, Figure S5). These findings imply two scenarios: (i) Sox2 is only partially saturated and undergoes intermediate-to-fast exchange between the free and DNA-bound state on the NMR timescale and (ii) Sox2 adopts two or more distinct conformations that resemble the free state while associated with weaker binding sites on nucleosomes. Since these spectra resemble the spectrum of Sox2 bound to a non-specific DNA at saturating concentrations (Figure S5), the second scenario is likely active. Titration of Oct4 to the Sox2-NCP 62_N_ sample, which based on EMSA results is expected to form is a stable ternary complex, shifted the resonance peaks closer to the DNA-bound state. This pointed to an increased population of the NCP-bound state of Sox2. However, even addition of excess Oct4 and nucleosome did not fully reproduce the spectrum observed for Sox2-DNA or -NCP 54R_F_ and left some peaks severely broadened, indicating persistent dynamics. These results suggest that Oct4-induced stabilization of the Sox2-NCP 62_N_ complex is not sufficient to mimic the stable complex of Sox2 with NCP 54R_F_ or with naked DNA.

### Impaired Sox2 folding and DNA bending activity inhibit nucleosome binding and Oct4 synergism

In the cryo-EM structure of Sox2 and Oct4 with a nucleosome resembling NCP 54R_N_, Sox2 binding entails disruption of the histone-DNA interface via a large DNA bending and insertion of the Sox2 C-terminal tail between the DNA and histone core.^17^ Interestingly, the Sox2 HMG domain N- and C-terminal tails are largely disordered in the free state^58^ and only become folded upon DNA binding and bending, forming interactions in the DNA minor groove and with one another and helix α3 (Figure 5A).^42, 44^ Thus, we wondered how altering the ability of Sox2 to bend DNA, through mutations in the protein N- and C-terminal tails, affects its ability to bind and bend nucleosomal DNA. Towards this, we engineered several single-point alanine mutations in the N-terminal (R7A), C-terminal (Y72A and Y74A), and α1-helix (N10A) of the Sox2 HMG domain (Figure 5A). R7 makes cross-strand hydrogen bonds with the nucleobase of T_6_(O2) and C_−5_(O2) in the minor groove of the mFgf4 motif and stacks against Y74, which in turn hydrogen bonds to the sugar of C_−5_ and the nucleobase of A_−6_(N3). Y72 does not directly interact with the DNA but is part of a hydrophobic core of residues (V5, H65, H69, and Y72) that stabilizes the C-terminal tail in the folded DNA-bound state. N10 inserts in the minor groove between residues T_4_ and G_5_, forming hydrogen bonds with the two nucleobases and thymine sugar. Since mutation at that position has been shown to reduce DNA bending,^59^ N10A was used as a control.

**Figure 5.**
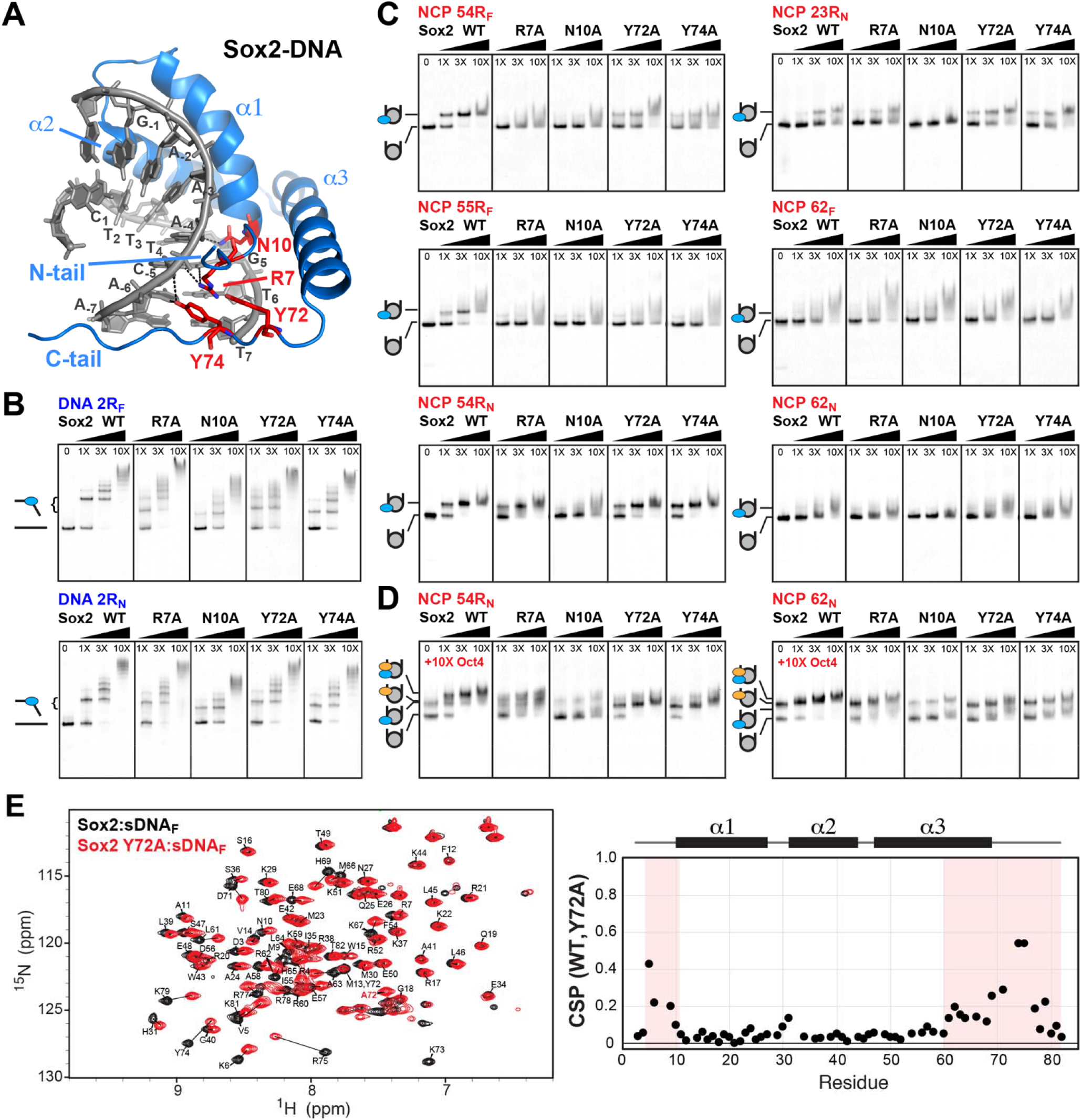
Impaired Sox2 folding and DNA bending activity differentially inhibit nucleosome binding and Sox2-Oct4 synergism. (A) Structure of Sox2 HMG domain (blue) bound to DNA (grey) containing the mFgf4 motif (numbered) (PDB ID: 1GT0). Highlighted in red are residues in the N-terminal tail (N-tail, R7), C-terminal tail (C-tail, Y72A and Y74A) or internal (α1-helix, N10) positions mutated in this study. (B) EMSA of Sox2 WT and mutants (0-100nM) binding to DNA (10nM) containing the mFgf4 (2R_F_) or Nanog (2R_N_) motif. The variable gel migration for the specific Sox2-DNA complex with different motif sequences or Sox2 mutations indicates variations in DNA bending (slower migration correlates with larger bending). (C) EMSA of Sox2 WT and mutants (0-100nM) binding to various NCP constructs (10 nM) showing overall larger effects of Sox2 mutations on the mFgf4 than mNanog motifs. (D) EMSA of Sox2 WT and mutants (0-100nM) binding to NCP 54R_N_ and 62_N_ (10nM) in the presence of Oct4 (100nM) showing that Sox2 mutants significantly impair the tertiary complex. (E) Overlay of ^1^H,^15^N-HSQC spectra of backbone amides for Sox2 WT (black) and Y72A (red) bound to a short DNA duplex (sDNA_F_) containing the mFgf4 motif. Chemical shift perturbations (CSPs) for Sox2 WT versus Y72A are plotted on the right as a function of residue number. Large CSPs (highlighted) and chemical exchange broadening are detected around the mutation site in the C-terminal tail and α3-helix, the N-terminal tail, and some internal residues, indicating changes in the conformation and dynamics of Sox2. Certain overlapped and severely broadened peaks as well as the mutated residue were omitted.

We first tested the interaction of Sox2 mutants with different nucleosome and DNA constructs using EMSA (Figure 5). Our data shows that binding of these mutants to DNA alters the electrophoretic mobility of the protein-DNA complex to a variable extent depending on the mutant identity and position of the Sox2 motif (Figure 5B). In general, migration of the mutant complexes was faster than that of the wild-type (WT) complex in the order N10A > Y74A ~ R7A > Y72A > WT for mFgf4 and N10A > Y74A > Y72A ~ R7A > WT for mNanog. This was evident when the Sox2 motif was located in the center of the DNA sequence (DNA 2R_F_ and 2R_N_, Figure 5B). DNA bending is expected to yield slower overall diffusion at the center rather than ends of the long duplex. Interestingly, binding of WT Sox2 appeared to generate an overall smaller bent for the mNanog than the mFgf4 site (Figure 5B). Moreover, the R7A mutant impaired DNA bending to a lesser extent at the mNanog than mFgf4 site, although the base-pairs that interact directly with the arginine sidechain are unchanged. The apparent DNA binding affinities for most Sox2 mutants were not substantially affected (K_d_ ~ 3.3 ± 0.5 nM for WT versus 4.4 ± 0.8 nM for R7A, 3.5 ± 0.6 nM for Y72A, and 5.6 ± 1.1 nM for Y74A for mFgf4), except for N10A (K_d_ ~ 19 ± 4 nM). However, the fraction of non-specific complexes was relatively increased in Sox2 mutants versus WT, indicating that the affinity for the cognate site was reduced.

For nucleosomes, we could not reliably assess the effect of Sox2 mutations or motif sequence on DNA bending. Association of Sox2 mutants with nucleosomes, however, showed distinct effects on the protein binding affinity and complex stability. For NCP 54R_F_ harboring the mFgf4 motif, the specific complex was largely abolished by R7A (K_d_ ~ 23 ± 3 nM) and, especially, N10A (K_d_ ~ 37 ± 5 nM) and visibly destabilized by the Y72A (K_d_ ~ 9.1 ± 1.0 nM) and Y74A (K_d_ ~ 5.5 ± 0.7 nM) mutants (Figure 5C). The effect of Sox2 mutations was even more dramatic when the mFgf4 motif was shifted by one base-pair in NCP 55R_F_ and a discrete band for the specific complex could no longer be detected (Figure 5C). By contrast, Sox2 mutants had a lesser impact on binding to the mNanog motif on nucleosomes (Figure 5C). For NCP 54R_N_, the specific complex could still be observed with all mutants (except for N10A), with small differences for R7A and Y72A. The impact of these mutations was less pronounced for the more internal position near SHL2 in NCP 23R_N_. In addition, weaker and non-specific binding was not substantially affected by the mutations. This data indicates that Sox2 mutations that reduce DNA bending, presumably by loss of direct contacts and/or proper Sox2 folding, also decrease the stability of the Sox2-nucleosome complex in a sequence-dependent manner. Moreover, the higher impact of the R7A mutation on DNA bending with mFgf4 than mNanog correlates with a larger decrease in stability of mFgf4-containing nucleosome complex. This suggests that the ability of Sox2 to optimally bend DNA motifs is important for the formation of a stable nucleosome complex.

At the same time, we observed that the Sox2-Oct4-NCP ternary complex on Nanog motifs (NCP 54R_N_ and 62_N_) was significantly depleted by Sox2 mutants that displayed small changes in nucleosome binding affinity alone (Figure 5D). Importantly, none of these mutations are found at the interface between Sox2 HMG and Oct4 POU_S_ or involved in direct protein-protein contacts on the Nanog motif. Sox2-Oct4 synergism on naked DNA containing the mNanog motif was also negatively impacted by the mutants (Figure S6). As a result of extensive signal overlap and smearing, we could not precisely quantify the decrease in cooperativity on nucleosomes. It is possible that the Sox2 C-terminal tyrosine mutations destabilize the folded protein conformation and alter the orientation or dynamics of HMG α3-helix that forms interactions with POU_S_, which could affect the geometry at the Sox2-Oct4 interface.

To determine whether Sox2 folding is affected by mutations in C-terminal tyrosines, we assessed the effect of the Y72A mutant on the conformation and dynamics of Sox2 in complex with DNA by NMR. We observed large chemical shift perturbations and/or enhanced line broadening of protein backbone amides and side-chain methyl groups near the mutations cite on the C-terminal tail (i.e. H69-K79), α3-helix (i.e. L61-E68) and at the N-terminal tail contacting Y72 (i.e. V5, K6) (Figure 5E and Figure S7). These signified pronounced changes in the conformation and dynamics of the Sox2 mutant in complex with DNA. Thus, it is possible that the Y72A mutation reduces overall DNA bending by destabilizing the folded conformation of Sox2 on DNA. Together with our EMSA results, these findings establish that the impaired ability of Sox2 to properly fold and bend DNA could lead to an altered conformation on nucleosomes and negatively impact the stability of the ternary complex with Oct4.

Finally, to examine the ability of Sox2 mutants to produce a stable bent DNA conformation in the context of nucleosomes, we performed the APB crosslinking assay with NCP 54R_F_ H2B(S53C) and the mutant proteins (Figure S6). We detected significant DNA cleavage for R7A and N10A mutants, which was reduced by roughly 50% relative to the free NCP. Binding of Y72A and Y74A mutants resulted in considerably less DNA cleavage (< 20%). The addition of Oct4 did not significantly affect the cleavage profile in the presence of Sox2 WT or mutants. These findings could be explained by variations in the binding affinities across Sox2 mutants as well as differences in the degree of DNA bending at the Sox2 binding site. The enhanced cleavage with R7A and N10A mutants, which show weaker affinity and do not produce a stable nucleosome complex by EMSA, could result from a larger fraction of free NCP. It is unclear whether the unstable complex reflects faster dissociation kinetics with a bent DNA conformation similar to the WT complex or less significant DNA deformation or both. Conversely, the reduced cleavage with Y72A and Y74A mutants, which retain tighter binding and form the stable complex, correlates with a highly bent DNA conformation. Further studies probing the kinetics of Sox2 binding to nucleosomes and the conformation of Sox2 mutants are required to discern between different scenarios. Overall, our mutational and NMR studies point to a key role of Sox2 HMG domain folding and coupled DNA bending in establishing a stable complex and Oct4 interactions on nucleosomes.

## Discussion

It remains largely unclear how pTFs recognize nucleosomes and target only a subset of their specific motifs in silent chromatin *in vivo*. One possible mechanism is that they exhibit strong preferences for specific positions on the nucleosome. Thus, well-defined nucleosome positioning within the genome can help guide pTFs localization to their optimal binding sites. In this study, we find a strong dependence of the binding affinity and stable complex formation of Sox2 and Oct4 on the position and sequence of their cognate motifs on nucleosomes. We observe that motifs where the minor groove of the Sox2 site or the major groove of the Oct4 POU_S_ or POU_HD_ site faces outward and away from the histone core are generally favored. Our findings are consistent with previous experimental and *in silico* studies showing preferential Sox2 and Oct4 binding to solvent-exposed sequences on nucleosomes.^17, 18, 23, 34, 46^ This is expected due to the high energetic cost of dissociating large segments of nucleosomal DNA from the histone octamer to accommodate protein binding on the inner surface, particularly at more internal positions. Additionally, unfavorable electrostatics between the positively charged pTFs and histones as well as steric and geometric constraints would further disfavor inner facing binding schemes.

While solvent accessibility plays a major role, it is not the only factor that determines the binding preference of pTFs. We find that Sox2 favors strongly only certain solvent exposed nucleosome sites. We observe a super-stable Sox2-nucleosome complex near SHL5 (NCP 54R) and somewhat less stable ones near SHL2 (NCP 23R) and SHL6 (NCP 65R). For those positions, the DNA minor groove faces outward and the Sox2 N- and C-terminal tails are oriented away from the neighboring DNA gyre. Prior high-resolution studies have captured only the Sox2-Oct4-NCP ternary complex where the pTF binding site is positioned identically as in NCP 54R.^17^ Here, by using gel shift, nuclease footprinting, and sensitive NMR methyl TROSY experiments, we demonstrate that Sox2 alone binds NCP 54R with high specificity and with similar affinity and conformation to that when binding to naked DNA. The affinity declines by several fold if the motif is (i) shifted by even one base pair (NCP 55R), (ii) translated to a similar rotational position externally to SHL6 (NCP 65R) or internally to SHL2 (NCP 23R) or (iii) reversed at the same location (NCP 52). By contrast, locations that have the Sox2 motif near the dyad, other internal positions, and certain entry/exit DNA sites in either orientation display considerably weaker affinity and, as detected by NMR, potentially alternate Sox2 conformations. Smearing of the bands observed by EMSA as well as peak broadening observed by NMR suggest that those complexes are more dynamic and may exhibit faster dissociation kinetics than complexes at optimal positions. Future quantitative studies examining the site-specific binding rates and conformational dynamics of Sox2 would shed light on kinetic and thermodynamic factors that influence nucleosome recognition.

Notably, we do not observe a stable Sox2-nucleosome complex at the dyad or multiple other solvent accessible positions. Our findings contradict previous studies showing preferential binding of Sox2 and other HMG domain proteins near the nucleosome dyad.^28^ The discrepancy could be due to non-specific Sox2 complexes at the dyad, observed here and by MD simulations,^46^ which do not generate a dramatic DNA bend and may be more abundant in the context of different (not 601) underlying sequences.^28^ Indeed, we find that even non-specific association of Sox2 with nucleosomes exhibits high apparent binding affinity (K_d_ ~ 25-50 nM), which is several fold higher than the specific binding of Oct4 to nucleosomes. We cannot exclude that this results from a strong non-cognate Sox2 binding site on the control 601* sequence in the context of nucleosomes. In that case, fast Sox2 dissociation rates may preclude observation of a distinct DNaseI footprint with the control NCP as the one observed with non-specific binding to the control DNA.

Why are many accessible nucleosome positions, including the dyad, disfavored by Sox2? The stability of the Sox2-nucleosome complex can be largely modulated by differences in the energetic penalty for DNA bending and disruption of histone-DNA contacts to accommodate Sox2. Structural analysis of nucleosomes shows that the DNA minor groove width and extent of helical deformation vary substantially among different sites with the same rotational setting.^60, 61^ Additionally, mapping of position-specific histone-DNA interactions by single-molecule DNA unzipping experiments^62^ and steered MD simulations of DNA unwrapping^63^ point to sizeable variations across the nucleosome. Even though the dyad has a single DNA gyre and allows for higher minor groove accessibility, it features stronger histone-DNA interactions^62, 63^ and smaller minor groove expansion^61^ than other positions. These factors and the central location of the dyad would disfavor large DNA deformations and long-range helical shifts that may accompany Sox2 binding. Similar considerations apply to other internal positions that maintain strong histone-DNA contacts. ^62, 63^ Conversely, it may seem surprising that we observe diminished binding near the DNA entry/exit points (SHL6 to SHL7), where spontaneous DNA unwrapping occurs. However, MD simulations indicate that protein-DNA contacts are actually enhanced near SHL7 relative to the region spanning SHL5 to SHL6, which is primarily due to histone H3 N-terminal and H2A C-terminal tail contacts with entry/exit DNA.^63^ Such interactions could make it more difficult to distort and displace the DNA helix at the edge of nucleosomes. Both NMR studies and MD simulations have uncovered extensive dynamic interactions between histone H3 tails and entry/exit DNA, which can impact the binding of other chromatin factors at the DNA entry/exit points.^64^ In that respect, it would be important to investigate the impact of targeted histone tail truncations or modifications on the binding of Sox2 to nucleosomes.

A main finding in our study is the strong preferential binding of Sox2 near SHL5, which mimics the pTF complex with naked DNA. The ability of Sox2 to strongly bind and bend DNA at that position, displacing the helix from the histone core, could stem from the overall weaker contacts between DNA and histone globular domains near SHL5.^63^ Examination of nucleosome structures shows that the inner facing minor groove near the Sox2 binding site is contacted by Arginine (Arg) or Lysine (Lys) residues only from the flexible H2A (SHL±4.5) and H2B (SHL±5) N-terminal tails.^61^ By contrast, all other inner minor groove positions interact with at least one Arg belonging to the globular histone domains.^61^ Recognition of the compressed minor groove of nucleosomal DNA (especially AT-rich sequences) by long and positively charged Arg and Lys side chains is critical for nucleosome stability and underlies the strong nucleosome positioning potential of the 601 sequence.^65^ Because of the flexible nature of histone tails,^64^ Sox2 binding near SHL5 may not entirely abolish DNA interactions with the histone tail Arg/Lys residues but instead relocate them to nearby grooves on the same (H2A) and/or adjacent (H2B) DNA gyre. Additionally, while Sox2 binding at that site causes large DNA deformation, as demonstrated by our crosslinking data and the published Sox2-Oct4-NCP structure,^17^ it does not unwrap or change substantially the conformation and spatial orientation of entry/exit DNA.^17^ This suggests that the nucleosome can accommodate Sox2 binding at SHL5 without large-scale DNA distortion and still retain critical contacts between histones and DNA at the edge of nucleosomes. Collectively, these factors could facilitate the preferential binding of Sox2 near SHL5. Alternatively, binding of Sox2 at a similar rotational setting near SHL6 or SHL2 is expected to disrupt histone core Arg/Lys contacts and directly displace histone H3/H2A (SHL6) or H4 (SHL2) tails. Therefore, the reduced Sox2 binding there as compared to SHL5 could be explained by a combination of stronger and more extensive histone contacts with DNA, which may not be easily re-established at nearby DNA.

The strong Sox2 binding that we observe near SHL2 could be aided by other factors such as DNA shape and sequence. The pre-bent conformation of DNA in nucleosomes, which creates wider outward facing minor grooves, aligns with the direction of Sox2-induced DNA bending and could pre-dispose the association of Sox2 (and other HMG proteins) with nucleosomes. Based on analysis of diverse nucleosome structures, the SHL2 position is generally characterized with the widest minor groove, on average ~2 Å wider than at the dyad.^60, 61^ This, coupled with somewhat weaker interactions between the globular histone domains and DNA at SHL2 relative to most other locations,^17^ could contribute to the enhanced binding at that site. A recent high-resolution study showed that Sox2 binding at a similar position near SHL2 leads to partial unwrapping of DNA on the adjacent gyre and enhanced dynamics of entry/exit DNA.^18^ This could provide a mechanism whereby Sox2 binding stimulates DNA opening and initial destabilization of nucleosomes. Currently, it is unclear whether this phenomenon is common in nucleosomes or whether the *in vitro* selected DNA sequence in that study is inherently more flexible and prone to dissociation than the 601 or other sequences. Indeed, higher structural flexibility have been observed by MD simulations for nucleosomes containing “natural” DNA sequences derived from gene regulatory regions featuring one or more pTF binding sites.^47^ This could be resolved by future experiments investigating the impact of DNA sequence on Sox2-induced perturbations in nucleosomes.

We also find that the sequence of the Sox2 recognition motif plays a key role in the stability of its complex with the nucleosome. The mFgf4 motif does not form a stable complex with Sox2 near SHL2, while the mNanog motif produces a relatively stable nucleosome complex and higher apparent affinity. Conversely, at SHL5, the mNanog motif yields lower affinity for Sox2 than the mFgf4 motif. Importantly, we uncover that Sox2 association with the mNanog motif on DNA generates smaller helical bending than with the mFgf4 motif. Since differences in the motif sequence are not expected to significantly alter specific protein-DNA interactions, this likely stems from inherent variations in DNA deformability. Sox2 binding to DNA produces a large kink from intercalation of a methionine residue at the central TpG dinucleotide step.^42^ It is plausible that sandwiching the central TpG step with two rigid A-tract^66^ segments (5’TTT and 5’TTTT, Figure 1A) in the mFgf4 sequence produces a greater DNA bent than in the mNanog sequence that lacks A-tracts (Figure 1A). Larger DNA bending of the mFgf4 sequence could, in turn, optimize Sox2 folding on DNA and thus offset the higher energetic cost of DNA deformation. This is in line with our findings that Sox2 mutations that compromise its DNA bending activity, by destabilizing protein folding (Y72A by NMR), also inhibit more strongly Sox2 binding to the mFgf4 than the mNanog motif. Together, these results imply that larger DNA deformation is required for specific recognition of mFgf4 than mNanog Sox2 motifs on nucleosomes. Moreover, it seems that distinct factors dominate Sox2 binding at different nucleosome positions. We propose that, at SHL5, disruption of histone-DNA contacts has a lower energetic cost and, thus, the direct (sequence) and indirect (shape) readout of nucleosomal DNA by Sox2 resembles more closely that of naked DNA. By contrast, at SHL2, DNA bending and dissociation from the histone core has a higher energetic penalty and, thus, Sox2 favors creation of the less bent complex on the mNanog motif. A smaller Sox2-induced DNA deformation at the SHL2 is consistent with cryo-EM structures of Sox2 bound at SHL2 (5’CTTTGTG)^18^ as compared to SHL5 (5’CTTTGTT)^17^ as well as with recent MD simulations.^67^ This model further implies that suboptimal Sox2 binding sites on naked DNA that generate smaller bending could be preferentially bound at certain nucleosome positions.

Finally, it is possible that the strong Sox2-nucleosome complexes we observe here are further stabilized by site-specific interactions between Sox2 and histones. For example, the Sox2 HMG domain could potentially interact with the histone H2A-H2B dimer near SHL5, particularly with the flexible H2A and H2B N-terminal tails that are found in the vicinity of SHL4 and SHL5, respectively. Also, the H4 and H2B N-terminal tails can transiently interact with DNA near SHL2, while the H3 N-terminal and H2A C-terminal tails localize at nucleosome edges near SHL6. Future experiments probing direct interaction between Sox2 and histones and investigating the effect of targeted manipulation of histone-DNA contacts could shed light into the positional preference of Sox2 on nucleosomes.

We further reveal that the Sox2 binding partner, Oct4, also exhibits strong positional preference for the major groove near nucleosome ends (SHL5.5 and SHL6.5) and mostly non-specific binding elsewhere, consistent with previous studies.^17, 46, 53^ The high-affinity complexes feature solvent exposed half-sites for either POU_S_ or POU_HD_. In the canonical Sox2-Oct4 complex on naked DNA, POU_S_ and POU_HD_ engage with opposite faces of the DNA helix (see Figure 1B).^42, 44^ This configuration is incompatible with nucleosome structure (see Figure 2C). Therefore, it is likely that one Oct4 subunit recognizes its specific site, while the other binds non-specifically to other nucleosome components or samples alternative conformations. This scenario is in line with the absence of electron density for the POU_HD_ subunit in cryo-EM structures^17^ and with recent MD simulations.^46, 47, 68^ Such flexibility and conformational exploration could be assisted by the long unstructured linker between the two Oct4 modules. The POU_HD_ accessible site (NCP 55R_F_), found only several base-pairs away from the entry/exit points, may be favored due to increased DNA propensity for unwrapping that could enable transient sampling of the POU_S_-DNA bound state. The POU_S_ accessible site is more internal and recognized by Oct4 with similar affinity in either orientation (NCP 54R_N_ and NCP 62_N_). This hints at an intrinsic preference for that position that could stem from optimal DNA shape or potential histone interactions. Since the truncated POU_S_ or POU_HD_ domains alone do not reproduce the binding profile of the full Oct4 DBD, interactions of the flexible linker and subunit with histones or nearby DNA may contribute to the binding affinity and specificity. This is supported by MD simulations showing that the Oct4 DBD adopts diverse configurations on nucleosomes engaging with both DNA and histones.^46, 47, 68^ Specifically, depending on the protein orientation, two distinct modes were captured *in silico* when POU_HD_ was bound near SHL5.5: one where POU_S_ interacts with DNA from the adjacent gyre near SHL2 and another where POU_S_ contacts the acidic patch of the histone H2A-H2B dimer.^46^ Such alternative binding modes could explain our experimental findings that Oct4 largely favors nucleosome end positions.

Moreover, we find that the synergism of Sox2 and Oct4 is highly context dependent, with both nucleosome position and spacing between motifs playing a role. The relatively weak binding of Sox2 to the mNanog motif at SHL6 (NCP 62_N_) is greatly enhanced in the presence of Oct4. This is demonstrated by the discrete ternary complex observed by EMSA and stabilization of the Sox2 bound state observed by NMR. There, Oct4 alone associates with high affinity and the POU_S_ site is optimally accessible and adjacent to the Sox2 HMG site, allowing for unobstructed protein-protein interaction. As seen in the corresponding cryo-EM structure,^17^ this complex shows a large Sox2-induced DNA bending, which profoundly alters the orientation of the linker DNA and may disrupt its interaction with histone tails (especially H3). While Oct4 alone binds stably at that position, the presence of Sox2 would be necessary to elicit the large DNA bending. This could have impact on inter-nucleosome packing or make histone tails more accessible for readout and enzymatic modification. By contrast, cooperative Sox2-Oct4 binding is scarce on the mFgf4 motif near SHL6 (NCP 62_F_), where the POU_S_ site is 3 base-pairs removed from the HMG site and occluded by histones. The minor ternary complex detected by EMSA could result from transient unwrapping of entry/exit DNA and accommodation of Oct4 as in the DNA-bound state or an alternative Oct4 conformation. The ternary complex could also be observed at the high-affinity Sox2 site near SHL5 and, likewise, only on the mNanog motif (NCP 54R_N_). Although we do not observe strong Oct4-mediated stimulation of Sox2 binding at that position, Sox2 folding and DNA bending could be potentially optimized via interactions with POU_S_. By contrast, Oct4 bound to NCP 54R_F_ and 55R_F_ containing the mFgf4 motif is displaced by excess Sox2, which indicates that Sox2 competes for the same site or allosterically inhibits Oct4 binding at a non-overlapping site. Importantly, the ternary complex is not observed at the dyad or internal positions containing the mNanog motif, including the higher-affinity Sox2 site at SHL2 (NCP 23R_N_). The presence of histones tails and more extensive histone-DNA interactions may preclude an optimal geometry of Sox2-Oct4 binding at more internal positions. In general, our findings agree with prior biochemical and single-molecule Forster resonance energy transfer (smFRET) studies showing that Sox2 and Oct4 bind cooperatively at end-positioned sites on nucleosomes but that this effect is attenuated or absent at the dyad and internal sites.^17, 69^ However, it is plausible that Sox2-Oct4 synergism is augmented in the context of nucleosomes that incorporate less positioning DNA sequences or destabilizing histone mutants and variants.

Lastly, we observe that Sox2-nucleosome interactions and synergism with Oct4 can be inhibited by mutations that impair the folding of the Sox2 N- and C-terminal tail, which is coupled to DNA binding. These findings provide mechanistic insight into how the folding and stability of the Sox2 HMG domain affects its ability to bind and deform DNA on nucleosomes. We further propose that, because of the prohibitively high energetic cost for DNA bending, the Sox2 HMG domain is unable to fold properly at most internal positions and establish a stable complex (on the consensus motif). Indeed, for more weakly binding NCP constructs, we observe by NMR that multiple Sox2 signals are shifted towards the free state or are severely broadened, similar to non-specific DNA binding. These findings are consistent with Sox2 sampling different conformations while bound to nucleosomes. However, since Sox2 folding and DNA bending depend on the DNA sequence, certain motifs might still be favored at internal positions (similar to mNanog at SHL2). Improper folding or conformational dynamics in the C-terminal tail and α3 helix, which we observe by NMR for the Y72A mutation, may also inhibit direct interactions with Oct4 POU_S_. Importantly, this could have broader implications for the ability of other HMG-containing proteins to bind the nucleosome and synergize with co-factors.

In summary, our findings demonstrate that pTFs Sox2 and Oct4 have optimal binding modes to nucleosomes that coincide with distinct solvent accessible positions and DNA sequences. The high-affinity nucleosome binding modes of Sox2 and Oct4 may be realized by optimal nucleosome positioning and guide pTF site selection in the cell. They might be further modulated by nucleosome composition (i.e. histone variants and modifications) and co-factor binding for more targeted chromatin localization and different functions. Extensive non-specific nucleosome binding, especially for Sox2, could also enhance pTFs chromatin association even when their cognate motifs are occluded or absent. Moreover, these diverse binding modes could facilitate the engagement of other TFs or affect the activity of chromatin enzymes and, thereby, promote chromatin opening and transcription.

## Materials and Methods

### Protein constructs and preparation

The human Sox2 HMG DNA-binding domain (39-118) was amplified by PCR from a modified pSpeedET plasmid^70^ and inserted into a modified pET15 plasmid^71^ containing an N-terminal (6X)-Histidine (His) tag and a 3C protease cleavage site using standard protocols. Sox2 single alanine mutations (R7A, N10A, F54A, Y72A, Y74A) were generated using standard mutagenesis protocols. The Oct4 DBD (133-296), POU_S_ domain (133-228), and POU_HD_ domain (229-296) were similarly inserted into a modified pET15 plasmid^70, 71^ containing an N-terminal (8X)-His tag, followed by an MBP tag and 3C protease cleavage site. All original plasmids were obtained from the DNASU plasmid repository and the 3C protease site was inserted in house. All DNA oligonucleotides used for plasmid modification and mutagenesis were obtained from IDT, Inc. (Coralville, IA). For large-scale protein expression, plasmids were transformed into BL21(DE3) cells and grown in TB medium with 100 mg/L carbenicillin (for non-isotopically labeled protein) at 37°C with shaking until a density of OD_600_ ~ 0.6. Protein expression was induced with 0.5 mM IPTG and cells grown for 4 hours at 37°C for Sox2 and 16-18 hours at 18°C for Oct4. For uniform ^15^N or ^15^N, ^13^C isotopic labeling, cells were grown in M9 Minimal media supplied with ^15^N-NH_4_Cl (1g/L) and ^13^C-D-glucose (2g/L). For unform ^2^H,^15^N-labeling with site-specific Ileδ1-[^13^CH_3_], Leu,Val-[^13^CH_3_, ^12^CD_3_], Ala-[^13^CH_3_] (ILVA) methyl labeling,^72^ the expression medium was prepared with ^15^N-NH_4_Cl_2_ (1g/L), ^2^H-D-glucose (3g/L), 99% D_2_O and the following labeled substrates were added 1 hour prior to induction: 2-^2^H,3-^13^C-L-alanine (0.7 g/L), ^2^H-succinate (2.5g/L), 3,4,4,4-^2^H,3-^13^CH_3_-α-ketoisovalerate (0.12g/L), and 3,3-^2^H,^13^CH_3_-α-ketobuterate (0.06g/L). Cells were acclimated to the high D_2_O content in the expression media by a gradual increase in the D_2_O/H_2_O ratio, as previously described.^72^ All isotopically labeled compounds were obtained from CIL. Cells were harvested by centrifugation at 4000rpm for 20 min at 4°C and pellets stored at −20°C until needed. For purification, cell pellets were resuspended in ~ 40ml (per 1L growth) of HisBind A buffer (50mM Tris pH 7.5, 500mM NaCl, 10mM Imidazole, 10% Glycerol, 5mM β-Mercaptoethanol (BME)) supplemented with 0.2mM PMSF, 1mM Benzamidine, 0.15mg/ml Lysozyme, and 1mM DTT, incubated on ice for 30 min and sonicated for 4-6 cycles of 60 sec on ice followed by centrifugation at 23,000g for 20 min at 4°C. The protein was purified by a two-step Ni^2+^ affinity chromatography on an AKTA FPLC system (GE Amersham Pharmacia). Specifically, the lysate was loaded on a 2×5ml HisTrap HP column (GE Healthcare) pre-equilibrated with HisBind A, washed for ~10 column volumes, and eluted with a linear gradient of buffer HisBind B (50mM Tris pH 7.5, 500mM NaCl, 500mM Imidazole, 10% Glycerol, 5mM BME) over 7-10 column volumes. Protein fractions were combined and diluted with 50mM Tris pH 7.5, 25mM NaCl, 1mM DTT to a final 130-150mM NaCl/~80mM Imidazole and the His tag was cleaved using an in-house (6X)His-tagged 3C protease (~ 1.5 mg/1L of growth) for 16 hours at 4°C. The His tag, 3C protease enzyme and impurities were removed by re-passing the protein on a 5ml HisTrap HP column pre-equilibrated with HisBind A buffer supplemented with 30mM Imidazole and eluted in the wash step. Protein purity was assessed by 18% SDS-PAGE and pure fractions were concentrated, flash frozen, and stored at −80°C.

*Xenopus Laevis* histones H2A, H2B, H3, and H4 in pET3a plasmid were a gift from Prof. Gregory Bowman (Johns Hopkins University) and were expressed and purified as previously outlined^73–75^ with some modifications. Briefly, histone plasmids were transformed in BL21(DE3) pLysS cells, grown in 2xTY medium with 100mg/L carbenicillin at 37°C with shaking to a density of OD_600_ ~ 0.6 and induced with 0.3mM IPTG for 4 hours at 37°C. Cells were harvested by centrifugation at 3000rpm for 20 min at 18°C and pellets stored at −20°C. Histones H2A, H2B and H3 (2L each) were purified using ion exchange chromatography under denaturing conditions according to a published method.^75^ Specifically, cell pellets were resuspended in buffer SAU 200 (40mM NaOAc pH 5.2, 7.8M Urea, 200mM NaCl, 1mM EDTA pH 8.0, 10mM lysine, 5mM BME) supplemented with protease inhibitors (0.2mM PMSF, 1mM Benzamidine) and lysed by sonication for 6 cycles of 60 sec, then spun down at 23,000g for 20 min at 4°C. The lysate was loaded on a 20ml tandem Q-SP HiTrap HP column (2×5ml Q on top of 2×5ml SP), washed for 10 column volumes, where the Q column was removed and the protein eluted from the SP column by a linear salt gradient over 10 col. vol. using buffer SAU 800 (40mM NaOAc pH 5.2, 7.8M Urea, 800mM NaCl, 1mM EDTA pH 8.0, 10mM lysine, 5mM BME). Protein fractions were analyzed by 18% SDS-PAGE, pooled and dialyzed 4 times against ddH_2_O with 5mM BME using a 3.5kDa cutoff membrane (Spectra Labs), then lyophilized and stored at −20°C. Histone H4 was prepared instead from inclusion bodies, followed by a desalting column (HiPrep 26/10 (GE)) and a tandem Q-SP column (GE), as previously described.^73, 74^

### Nucleosome constructs and preparation

The Sox2-Oct4 mFgf4 and mNanog composite sites were incorporated at different positions within the Widom 601 positioning sequence^50^ in pGEM-3z/601 plasmid (gift from Prof. Gregory Bowman, Johns Hopkins University), which was modified on one end to remove a strong non-cognate Sox2 binding site (see Table S1). Nucleosomal DNA was prepared by standard large-scale PCR (10-40ml) of the modified 601 sequences using in-house Taq DNA polymerase, 10X ThermoPol buffer (NEB), 0.5 mM dNTPs (Invitrogen) and either unlabeled DNA primers (for NMR) or fluorescently 5’-end labeled (5’-(6-FAM) on the top strand shown in Table S1, 5’-Cy3 on the bottom strand) primers obtained from IDT with HPLC purification (see Table S1). DNA was concentrated using an Amicon Ultra-15 centrifugal filter unit (10kDa cutoff, Millipore-Sigma) and purified by vertical gel electrophoresis on a 6% (60:1 Acrylamide/Bisacrylamide) native gel column using a 491 Prep Cell (Bio-rad) run at 10W and 4°C, as described.^74^ DNA fractions were analyzed on a 1.2% agarose gel, concentrated and stored at −20°C. DNA concentration was determined from absorbance at 260nm measured on a NanoDrop device (Thermo Scientific) and using the extinction coefficient for duplex DNA (40ug/A_260_). *Xenopus Laevis* histone octamer was assembled by refolding each histone in 20mM Tris-HCl pH 7.8, 6M Guanidine-HCl, 5mM DTT at 2mg/ml and combining the four histone proteins with 20% molar excess of H2A and H2B (1:1:1.2:1.2), dialyzing 3-4 times against high salt buffer (10mM Tris-HCl pH 7.8, 2M NaCl, 1mM EDTA, 5mM BME), followed by concentration to ~ 1.4ml and gel filtration chromatography using a HiLoad 16/600 Superdex 200 pg column (GE), as per established protocols.^73, 74^ The octamer and H2A-H2B dimer fractions were analyzed by 18-20% SDS-PAGE, concentrated and flash frozen at −80°C for storage. Nucleosomes were reconstituted by combining purified histone octamer (6uM) with H2A-H2B dimer (1.8uM) and slight excess of DNA (6.6uM) in RBHigh buffer (10mM Tris pH 7.8, 2M KCl, 1mM EDTA, 1mM DTT), followed by salt gradient dialysis with RBLow buffer (10mM Tris pH 7.8, 250mM KCl, 1mM EDTA, 1mM DTT) over ~24 hours at 4°C.^73, 74^ Nucleosomes were dialyzed overnight into elution buffer (10mM Tris pH 7.8, 2.5mM KCl, 1mM EDTA, 1mM DTT), concentrated, and purified by vertical gel electrophoresis on a 7% native gel column, as described for DNA above.^74^ Nucleosome fractions were analyzed using a 5% native gel and visualized by ethidium bromide staining or fluorescence imagining on a Typhoon 5 laser scanner (Amersham). Nucleosomes were concentrated, flash frozen (10% glycerol added), and stored at −80°C.

### Electrophoretic mobility shift assay (EMSA)

For EMSA, 10nM of 5’-FAM/5’-Cy3-labeled nucleosome (NCP) or DNA were mixed with variable concentrations of Sox2 or Oct4 protein (10nM to 1000nM) in binding buffer (10mM Tris-HCl pH 7.5, 150mM KCl, 1mM EDTA pH 8.0, 1mM DTT, 0.5mg/ml BSA, 5% glycerol) in a 20μl total reaction volume and incubated at 25°C for 30 min. Binding reactions (2μl) were loaded on a 5% native gel and run for 85 min (DNA) or 120 min (NCP) at 100V at 4°C. FAM and Cy3 fluorescence was measured on a Typhoon 9450 or 5 imager (GE Amersham) using excitation at 488 nm and 532 nm, respectively, and the appropriate emission filters (standard protocol). The signal (volumes) corresponding to free and bound NCP (or DNA) species were quantified using ImageJ and the fraction bound (F_Bound_) was calculated from the ratio of bound to total NCP (or DNA). F_Bound_ as a function of total protein concentration was fitted in Mathematica (Wolfram) to a 2-state binding model using the following quadratic equation to extract the apparent dissociation constant (see Table 1): F_bound_(NCP) = (([N] + K_d_+ [P]) – (([N] + K_d_ + [P])^2^ – 4[N][P])^1/2^)/(2[N]), where [N] is the total concentration of NCP or DNA (10nM), [P] is the total concentration of protein added (Sox2 or Oct4), and K_d_ is the apparent dissociation constant. The K_d_ values and errors reported in Table 1 are the average and standard deviation of at least three technical replicates.

### Dnase I footprinting assay

Binding reactions (25μl) of 100nM NCP or DNA with or without 1μM Sox2 and/or Oct4 were set up in 10mM Tris-HCl pH 7.5, 100mM KCl, 1mM DTT, 0.2mg/ml BSA, 0.02% Tween-20, 8% Glycerol and incubated at 25°C for 30 min. Next, 15μl of DnaseI solution, prepared from DNaseI enzyme and 10X DnaseI reaction buffer (NEB), was added to 25μl of binding reactions so that the final conditions contained DNase I at 0.0125 U/ul for NCP and 0.0025 U/ul for DNA as well as 2.5mM MgCl_2_, 0.5mM CaCl_2_, and 100mM KCl. The reactions were incubated for 5 min at 25°C and stopped by adding 40μl of quench buffer (10mM Tris-HCl pH 7.5, 50mM EDTA, 2% SDS, 300ng/ul Glycogen (Invitrogen)) and heating at 75°C for 30 min. To purify DNA, reactions were mixed with an equal volume (80μl) of 25:24:1 phenol:chloroform:isoamyl alcohol (Sigma), briefly vortexed and centrifuged for 2 min at full speed (15,000 rpm). The aqueous (top) layer was placed in a clean tube and 80μl of 24:1 chloroform:isoamyl alcohol was added, vortexed and spun down as above. The top layer (~80μl) was transferred to a clean tube again and DNA was ethanol precipitated overnight at −20°C by mixing in 3 vol. of 100% Ethanol, 0.1 vol. of 3M Sodium Acetate, pH 5.2, 0.04 vol. of 1M MgCl_2_, and 1μl of 5mg/ml Salmon Sperm DNA (Invitrogen). The mixture was centrifuged at full speed (15,000 rpm) for 30 min at 4°C and pellets further washed twice with 80% ice cold ethanol by spinning at 4°C for 10 min. The pellets were air dried for 30-60 min to remove any trace of ethanol and resuspended in 6μl of formamide loading buffer (89mM Tris-borate pH 8.3, 5mM EDTA, 95% formamide (deionized), 0.2% Orange G dye). The samples were directly run on an 8% denaturing sequencing gel (8% (19:1) Acrylamide/Bisacrylamide, 8M Urea, 1X TBE) at 65 W for 1.5 hours. Sequencing ladders were generated using the Thermo Sequenase Dye Primer Manual Cycle Sequencing Kit (Applied Biosystems). Gels were scanned within optically clear glass plates (VWR) and the FAM- or Cy3-labeled DNA fragments were visualized separately on a Typhoon 5 laser scanner (Amersham) as described for EMSA.

### Site-specific nucleosome crosslinking assay

An NCP 54R_F_ construct was prepared harboring a single Cysteine (C) mutation in H2B (S53C) found in close proximity to the DNA backbone in the Sox2 binding motif. The H2B(S53C) histone octamer was a gift from Prof. Gregory Bowman (Johns Hopkins University) and has been shown to effectively crosslink with nucleosomal DNA in nucleosome sliding studies.^56^ APB labeling of nucleosomes was performed following published protocols with some modifications.^55, 76^ Briefly, in the dark, the photolabile crosslinker 4-azidophenacyl bromide^77^ (APB, Abcam) was dissolved in 100% DMF to ~100mM and further diluted to 5mM in 10mM Tris-HCl pH 7.5, 10% Glycerol, 20% DMF. APB at 250μM final was added to 1μM NCP 54R_F_ H2B(S53C), which had been buffer exchanged into a 10mM Tris-HCl pH 7.5, 5% glycerol using an Amicon Ultra-0.5 (10kDa cutoff) centrifugal filter unit to remove DTT. The labeling reaction was incubated for 3 hours at 25°C and quenched by addition of 1M DTT to a final of 10mM. Binding reactions of 100nM NCP with 1μM Sox2 and/or Oct4 were set up in binding buffer as for EMSA and incubated for 30 min at 25°C. The reactions were irradiated with ultraviolet (UV) light for 1 min in the dark, mixed with 60μl of 20mM Tris-HCl pH 8.0, 0.2% SDS, 150mM NaCl, briefly vortexed, and heated at 70°C for 20min. An equal volume (~80μl) of 5:1 phenol:chloroform (Sigma) was added to the mixture, briefly vortexed, and spun down for 2 min at 13,000rpm, where the crosslinked DNA ends up near the interface of the organic and aqueous phase. The samples were further processed and purified as previously described.^76^ Briefly, crosslinked DNA was washed (4 times with 1.0M Tris-HCl pH 8.0, 5% SDS) and purified by ethanol precipitation at 4°C overnight, cleaved selectively at the crosslinked site by base-catalyzed hydrolysis by adding 0.1M NaOH for 40 min at 90°C, neutralized with HCl, and ethanol precipitated at −20°C overnight. The dried DNA pellets were resuspended in 6μl of formamide buffer and run on an 8% sequencing gel, scanned and processed as described above for DNaseI footprinting.

### NMR spectroscopy

NMR samples were prepared by exchanging the appropriate amounts of Sox2 protein or a mixture of Sox2 with unlabeled DNA, NCP 3 times into NMR buffer (20 mM Tris-HCl pH 7.0, 25 mM NaCl, 0.5 mM TCEP, 1 mM EDTA) using an Amicon Ultra-15 (10kDa cutoff) filter unit and centrifugation at 4000rpm and 4°C. Samples were adjusted to ~300μl and 7.5% D_2_O, placed in a Shigemi tube, and stored at 4°C for 0-2 days before NMR data collection. All NMR experiments were carried out on a Bruker Avance 600MHz or Ascend 800MHz NMR spectrometer equipped with a triple-resonance cryoprobe. Backbone and sidechain assignments for the Sox2 HMG domain (WT and Y72A), free or bound to a short DNA duplex containing the mFgf4 site (sDNA_F_, 5’ GACT<u>CTTTGT</u>TTGGC), were obtained from a standard suite of experiments (2D HSQC and 3D HNCO, HNCA, HNCACB, HNCOCACB, HCCH-TOCSY, and NOESY) acquired with non-uniform sampling (NUS) on a 0.4 mM ^13^C,^15^N-labeled protein sample at 25°C (~3 days collection time on 600MHz). Backbone amide chemical shift perturbations (CSPs) for the Sox2 Y72A versus WT complex with sDNA_F_ were calculated from the difference in proton (δ_H_) and nitrogen (δ_N_) chemical shifts in ^1^H,^15^N-HSQC spectra as follows: CSP = (0.1Δδ_N_^2^ + Δδ_N_^2^)^1/2^. ^1^H,^13^C-HMQC (methyl TROSY) correlation spectra were collected with 256 scans on 800MHz NMR at 25°C (~12 hours) on 20μM samples of ILVA-methyl labeled Sox2 free or bound to unlabeled NCP 62_F_, 62_N_, 54R_F_, 27_F_ or DNA 27_F_ in 1:1.1 or 1:2.1 molar ratio. To make the ternary complex of ILVA-labeled Sox2 with NCP 62_N_ and Oct4 at 1:2:1 and 1:2:2 ratios, unlabeled Oct4 DBD in NMR buffer was added to the Sox2-NCP (1:2) sample at 1 and 2 molar equivalents and the volume was adjusted down to 300μl by centrifugation at 10,000rpm and 4°C using an Amicon Ultra-0.5 (10kDa cutoff) filter unit. All NMR data was processed using nmrPipe^78^ and NESTA-NMR^79^ (NUS data), and analyzed using Sparky,^80^ Mathematica (Wolfram), and in-house scripts. Structural alignments and visualizations were performed using PyMOL (Schrödinger, LLC).

## Acknowledgements

We thank Prof. Gregory Bowman at Johns Hopkins University for kindly providing materials and access to instrumentation, stimulating discussions, and critical review of the manuscript. We also thank Dr. Samaneh Kondalaji (Washington University at Saint Louis) for providing critical comments on the manuscript. We are very thankful to members of the Bowman lab, particularly Dr. Ilana Nodelman, for help with various experimental procedures and many beneficial discussions. We are grateful to Dr. Ananya Majumdar, director of the Biomolecular NMR center at Johns Hopkins University, for assistance with NMR data collection and processing. This work was supported by start-up funds provided to E.N.N. by Johns Hopkins University.

## Notes

### Competing Interest Statement

The authors have declared no competing interest.

