## Supplementary Table 1 and Supplementary Figures 1 to 7 for "Nucleosome topology and DNA sequence modulate the engagement of pioneer factors SOX2 and OCT4"

*To whom correspondence should be addressed:

**Table S1.** Modified 601 DNA sequences used for naked DNA or nucleosome constructs in this study containing the Sox2-Oct4 composite motif (highlighted in cyan). The G-C base-pair in the Sox2 motif used for construct numbering is highlighted in magenta, the dyad position is indicated in bold, and the modified segment from the original Widom 601 sequence is underlined in DNA 601*. The top strand is only shown for clarity.

| Construct | DNA sequence |
|---|---|
| ***mFgf4*** | |
| DNA $2R_F$ | 5'TGGAGAATCCCGGTGCCGAGGCCGCTCAATTGGTCGTAGACAGCTCTAGCACCGCTTAAACGCACGTACGCT<br>**T**TGTTTGGATGCTAATAACCGCCAAGGGGATTACTCCCTAGTCTCCAGGCACGTGTGGTTACGAGGCTATCGT |
| DNA $23R_F$ | 5'TGGAGAATCCCGGTGCCGAGGCCGCTCAATTGGTCGTAGACAGCTCTAGCACCGCTTAAACGCACGTACGCG<br>**C**TGTCCCCCGCGTTTTAACCTTTGTTTGGATGCTAATCTAGTCTCCAGGCACGTGTGGTTACGAGGCTATCGT |
| DNA $54R_F$ | 5'TGGAGAATCCCGGTGCCGAGGCCGCTCAATTGGTCGTAGACAGCTCTAGCACCGCTTAAACGCACGTACGCG<br>**C**TGTCCCCCGCGTTTTAACCGCCAAGGGGATTACTCCCTAGTCTCCAGGCCTTTGTTTGGATGCTAATATCGT |
| DNA $55R_F$ | 5'TGGAGAATCCCGGTGCCGAGGCCGCTCAATTGGTCGTAGACAGCTCTAGCACCGCTTAAACGCACGTACGCG<br>**C**TGTCCCCCGCGTTTTAACCGCCAAGGGGATTACTCCCTAGTCTCCAGGCACTTTGTTTGGATGCTAATTCGT |
| DNA $65R_F$[a] | 5'TGGAGAATCCCGGTGCCGAGGCCGCTCAATTGGTCGTAGACAGCTCTAGCACCGCTTAAACGCACGTACGCG<br>**C**TGTCCCCCGCGTTTTAACCGCCAAGGGGATTACTCCCTAGTCTCCAGGCACGTGTGGTTACTTTGTTTGGA |
| DNA $0_F$ | 5'TGGAGAATCCCGGTGCCGAGGCCGCTCAATTGGTCGTAGACAGCTCTAGCACCGCTTAAATTAGCATCCAAA<br>**C**AAAGCCCCGCGTTTTAACCGCCAAGGGGATTACTCCCTAGTCTCCAGGCACGTGTGGTTACGAGGCTATCGT |
| DNA $27_F$ | 5'TGGAGAATCCCGGTGCCGAGGCCGCTCAATTGGTCGTAGACAGCTCTAGCACCGCTTAAACGCACGTACGCG<br>**C**TGTCCCCCGCGTTATTAGCATCCAAACAAAGACTCCCTAGTCTCCAGGCACGTGTGGTTACGAGGCTATCGT |
| DNA $32_F$ | 5'TGGAGAATCCCGGTGCCGAGGCCGCTCAATTGGTCGTAGACAGCTCTAGCACCGCTTAAACGCACGTACGCG<br>**C**TGTCCCCCGCGTTTTAACATTAGCATCCAAACAAAGCTAGTCTCCAGGCACGTGTGGTTACGAGGCTATCGT |
| DNA $52_F$ | 5'TGGAGAATCCCGGTGCCGAGGCCGCTCAATTGGTCGTAGACAGCTCTAGCACCGCTTAAACGCACGTACGCG<br>**C**TGTCCCCCGCGTTTTAACCGCCAAGGGGATTACTCCCTATTAGCATCCAAACAAAGGTTACGAGGCTATCGT |
| DNA $62_F$ | 5'TGGAGAATCCCGGTGCCGAGGCCGCTCAATTGGTCGTAGACAGCTCTAGCACCGCTTAAACGCACGTACGCG<br>**C**TGTCCCCCGCGTTTTAACCGCCAAGGGGATTACTCCCTAGTCTCCAGGATTAGCATCCAAACAAAGTATCGT |
| ***mNanog*** | |
| DNA $2R_N$ | 5'TGGAGAATCCCGGTGCCGAGGCCGCTCAATTGGTCGTAGACAGCTCTAGCACCGCTTAAACGCACGTACGCA<br>**T**TGTAATGCAAAATTTAACCGCCAAGGGGATTACTCCCTAGTCTCCAGGCACGTGTGGTTACGAGGCTATCGT |
| DNA $23R_N$ | 5'TGGAGAATCCCGGTGCCGAGGCCGCTCAATTGGTCGTAGACAGCTCTAGCACCGCTTAAACGCACGTACGCG<br>**C**TGTCCCCCGCGTTTTAACCATTGTAATGCAAAATCCCTAGTCTCCAGGCACGTGTGGTTACGAGGCTATCGT |
| DNA $54R_N$ | 5'TGGAGAATCCCGGTGCCGAGGCCGCTCAATTGGTCGTAGACAGCTCTAGCACCGCTTAAACGCACGTACGCG<br>**C**TGTCCCCCGCGTTTTAACCGCCAAGGGGATTACTCCCTAGTCTCCAGGCCATTGTAATGCAAAAGCTATCGT |
| DNA $27_N$ | 5'TGGAGAATCCCGGTGCCGAGGCCGCTCAATTGGTCGTAGACAGCTCTAGCACCGCTTAAACGCACGTACGCG<br>**C**TGTCCCCCGCGTTTATTTTGCATTACAATGACTCCCTAGTCTCCAGGCACGTGTGGTTACGAGGCTATCGT |
| DNA $62_N$ | 5'TGGAGAATCCCGGTGCCGAGGCCGCTCAATTGGTCGTAGACAGCTCTAGCACCGCTTAAACGCACGTACGCG<br>**C**TGTCCCCCGCGTTTTAACCGCCAAGGGGATTACTCCCTAGTCTCCAGGCACTTTTGCATTACAATGTATCGT |
| ***No motif*** | |
| DNA 601* | 5'TGGAGAATCCCGGTGCCGAGGCCGCTCAATTGGTCGTAGACAGCTCTAGCACCGCTTAAACGCACGTACGCG<br>**C**TGTCCCCCGCGTTTTAACCGCCAAGGGGATTACTCCCTAGTCTCCAGGCACGTGTGGTTACGAGGCTATCGT |

[a] DNA $65R_F$ contains a truncated composite motif that lacks the Oct4 binding site.

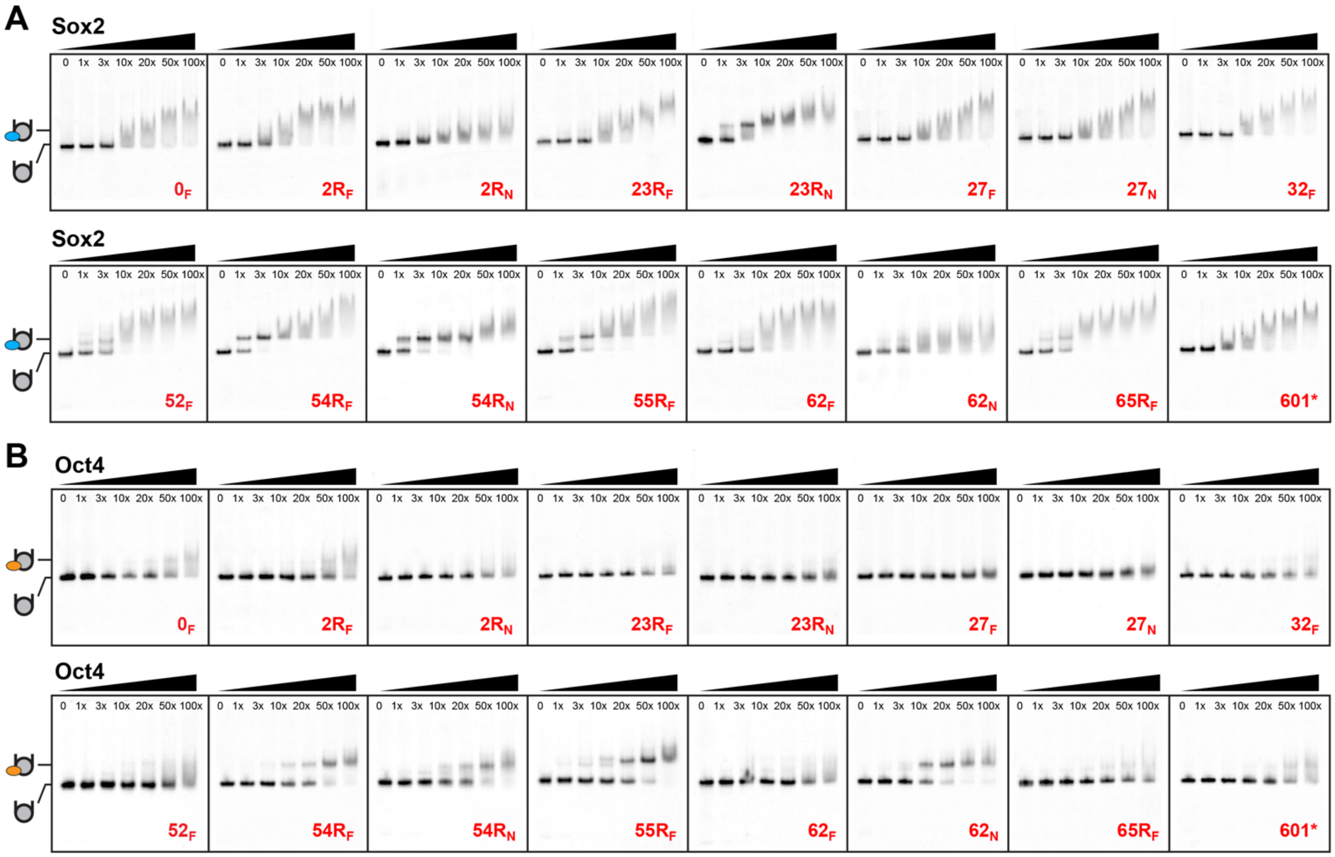


**Figure S1.** Pioneer factor Sox2 and Oct4 binding to nucleosomes is highly position and sequence dependent. Representative EMSA gels of (A) Sox2 and (B) Oct4 DBD binding titrations (0 to 1000 nM) to various nucleosome constructs (10 nM) containing the mFgf4 (subscript F) or mNanog (subscript N) composite motif. All binding reactions were visualized by FAM fluorescence.

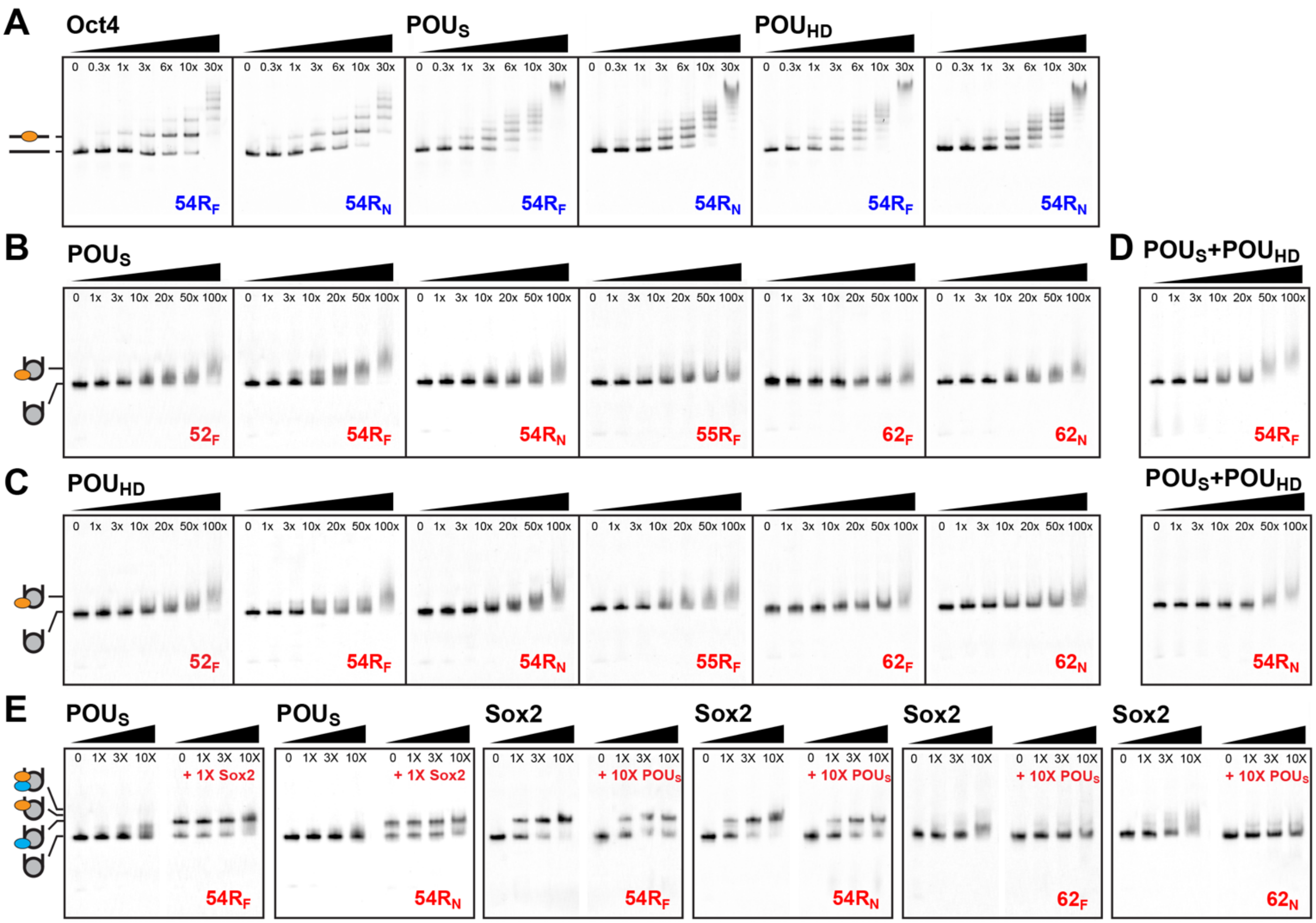


**Figure S2.** The full Oct4 DBD is required for efficient nucleosome binding alone and for cooperative binding with Sox2. (A) Representative EMSA gels of Oct4 DBD, $POU_S$, and $POU_{HD}$ binding to naked DNA (10 nM), indicating similar specific affinities and interaction at multiple sites. Representative EMSA gels for (B) $POU_S$, (C) $POU_{HD}$ or (D) $POU_S$ and $POU_{HD}$ proteins binding to various nucleosome constructs (10 nM) showing the formation of an unstable complex between either Oct4 subunits, alone or together, and NCPs where the full Oct4 DBD binds efficiently (NCP $54R_F$, $54R_N$, $55R_F$, $62_N$, Figure 2). (D) EMSA gels showing that the tertiary complex cannot be detected for Oct4 $POU_S$ in the presence of Sox2 and vice versa for NCPs, where a stable complex forms with the full Oct4 DBD (NCP $54R_N$ and $62_N$, Figure 2). All binding reactions were visualized by FAM fluorescence.

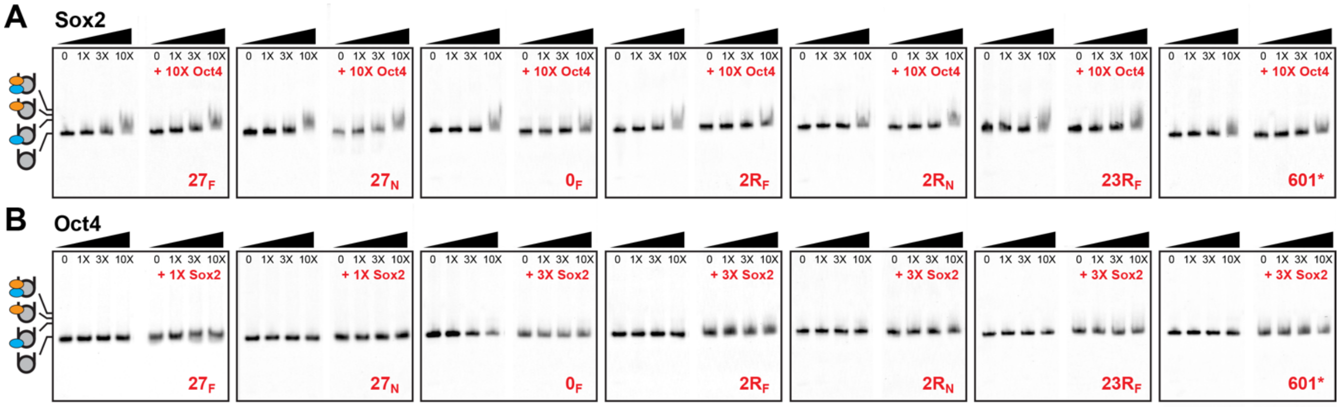


**Figure S3.** Sox2 and Oct4 do not visibly cooperate at internal nucleosome positions and different composite motif sequences. Representative EMSA gels of (A) Sox2 binding in the absence or presence of Oct4 and (B) Oct4 binding in the absence or presence of Sox2 showing the lack of stable tertiary complex formation for NCPs containing the composite motif at the dyad (NCP $0_F$ and $2R_F$), SHL2 to SHL3 (NCP $23R_F$, $27_F$, and $27_N$), and the control (NCP 601*). All binding reactions were visualized by FAM fluorescence.

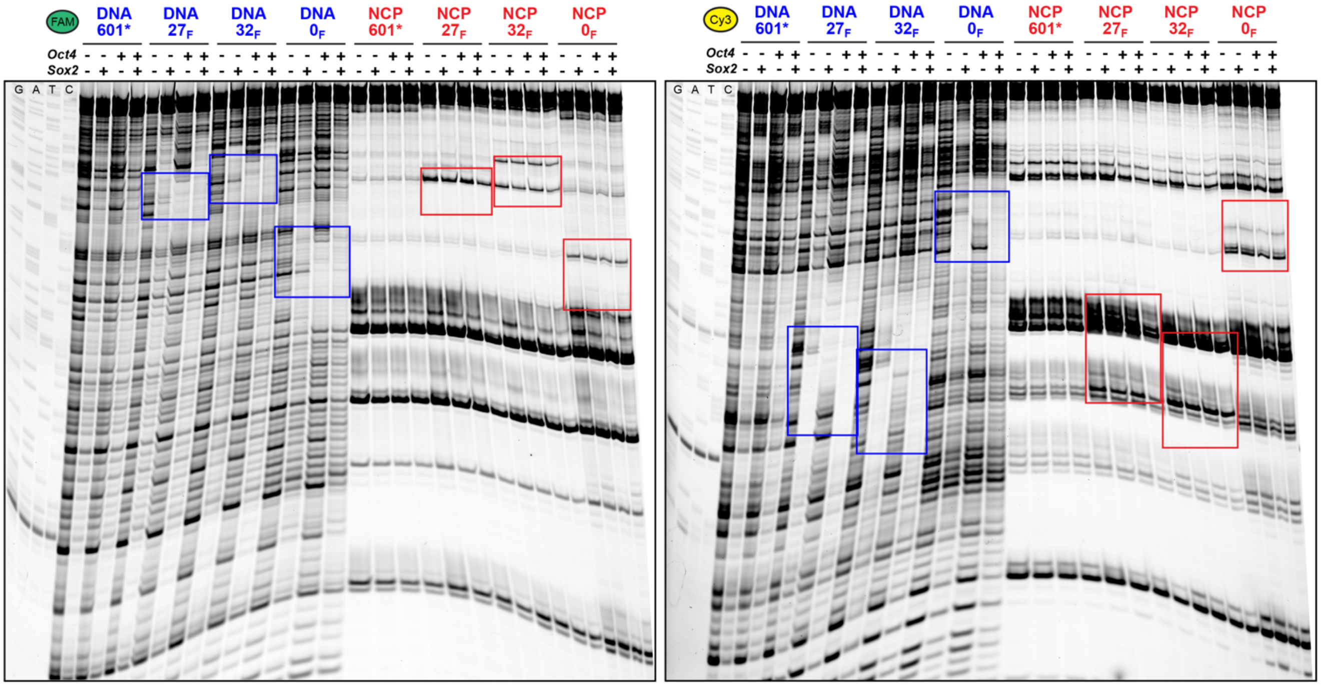


**Figure S4.** DNaseI footprinting of Sox2 and Oct4 binding to different DNA or NCP constructs. A clear footprint is observed for Sox2 and/or Oct4 binding to the composite motif (rectangles) on DNA, while no apparent footprint is observed for NCPs that exhibit weak protein binding by EMSA. The top strand (Table S1) is visualized by FAM fluorescence and the bottom strand is visualized by Cy3 fluorescence.

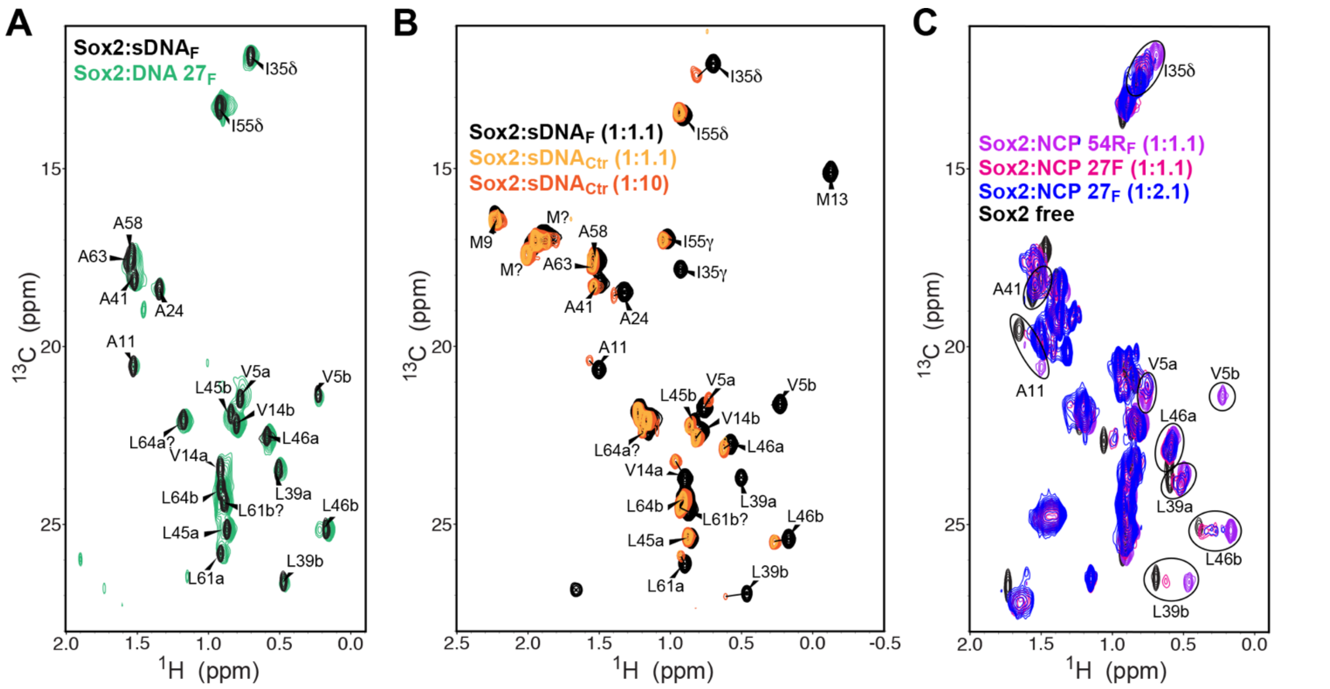


**Figure S5.** Sox2 binding to non-specific DNA resembles the conformation of the weaker nucleosome complex. (A) Overlay of methyl TROSY ($^{1}$H,$^{13}$C-HMQC) NMR spectra of the ILVA-labeled Sox2 HMG domain bound to specific short DNA (black, sDNA$_F$) and 601*-based long DNA (green, DNA 27$_F$) containing the mFgf4 site showing similar protein conformation. (B) Overlay of $^{1}$H,$^{13}$C-HSQC spectra of the $^{13}$C,$^{15}$N-labeled Sox2 HMG domain bound to a specific DNA (sDNA$_F$, black) in 1:1.1 ratio and non-specific DNA (sDNA$_{Ctr}$) in 1:1.1 (yellow) and 1:10 (orange) ratio. The spectra show chemical shift changes and enhanced chemical exchange (line broadening) in the non-specific complex, which persist at saturating DNA concentrations (1:10). (C) Overlay $^{1}$H,$^{13}$C-HMQC spectra of the ILVA-labeled Sox2 HMG domain free or bound to NCP 54R$_F$ (tight binder) and NCP 27$_F$ (weak binder) in 1:1.1 and 1:2.1 ratios, showing persistent chemical shift differences and chemical exchange (line broadening) in NCP 27$_F$ that resemble the behavior of the non-specific DNA complex in (B).

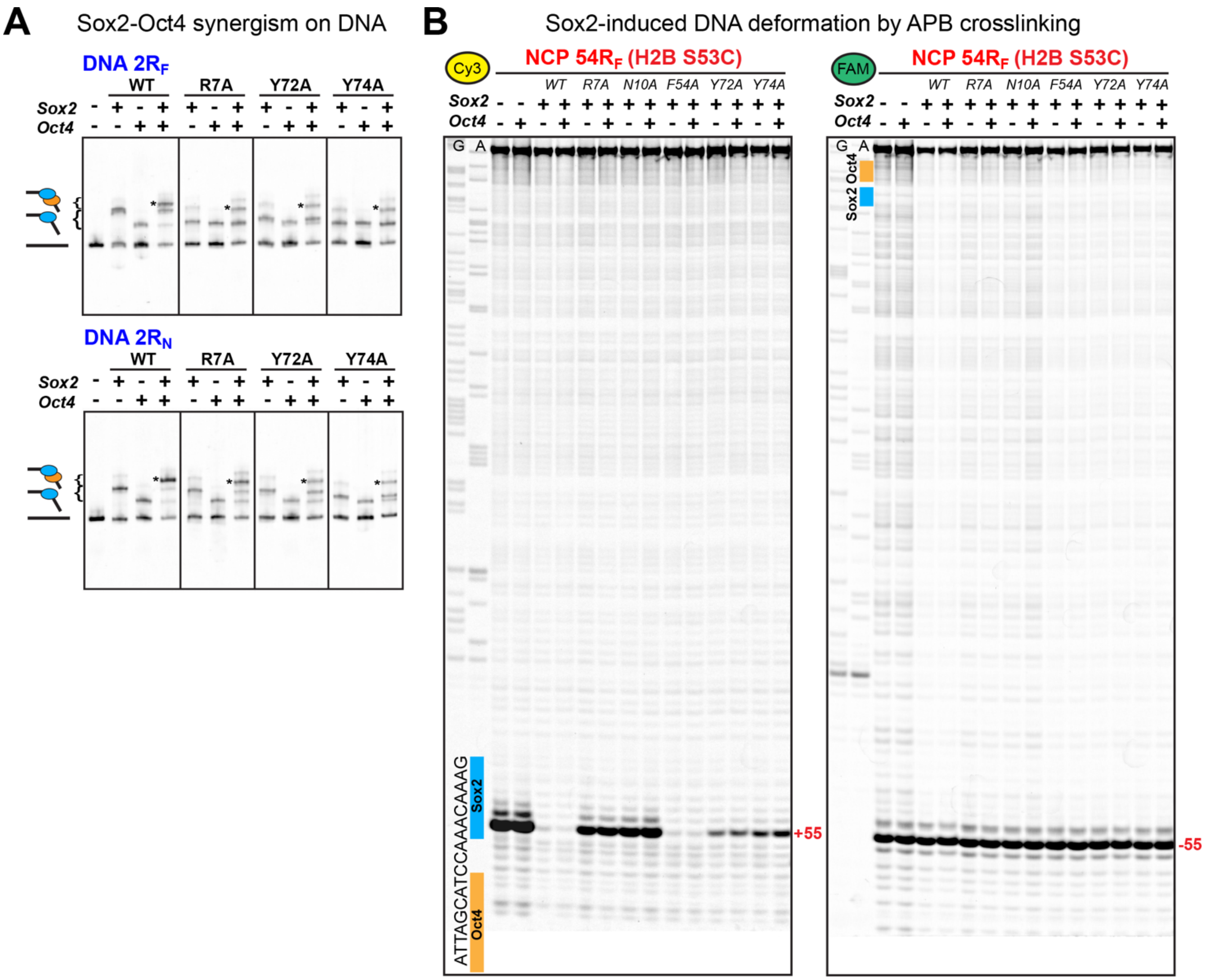


**Figure S6.** Sox2 mutations that impair DNA bending and protein folding inhibit binding and synergism with Oct4 on DNA and nucleosomes. (A) EMSA native gels showing that Sox2 N- and C-terminal tail mutations reduce DNA bending, as indicated by faster gel migration than the WT protein. The mutations also inhibit Sox2-Oct4 cooperativity, as indicated by the smaller fraction of the tertiary complex (*). (B) APB crosslinking assay of NCP 54$R_F$ H2B (S53C) in the presence of Sox2 and/or Oct4 showing differentially reduced DNA cleavage by Sox2 mutations, consistent with faster binding kinetics or reduced DNA deformation.

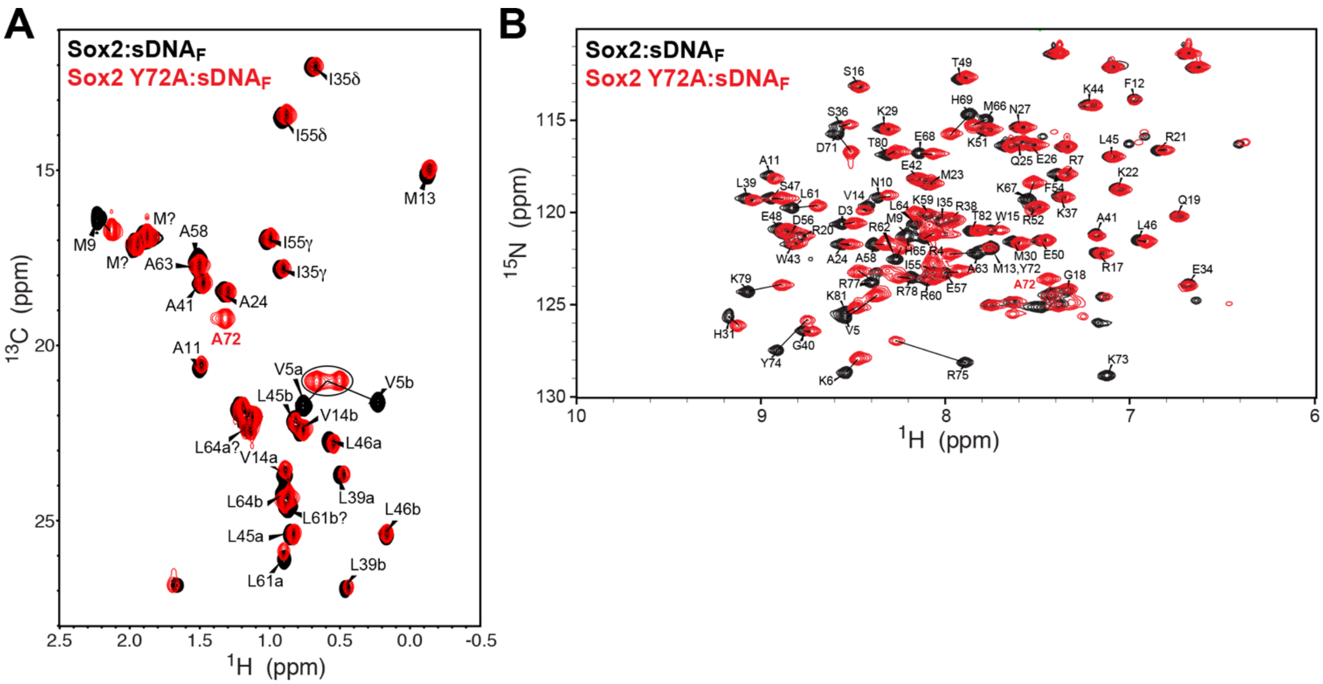


**Figure S7.** Sox2 C-terminal tail mutation affects the conformation and dynamics of the DNA bound state. Overlay of HSQC spectra of (A) sidechain methyls and (B) backbone amides of Sox2 wild-type (black) and Y72A mutant (red) bound to a short DNA duplex ($sDNA_F$) containing the mFgf4 motif. The data show large chemical shift perturbations and exchange (peak broadening) near the Y72A mutation site (i.e. H69 to K79), the N-terminal tail (i.e. V5, K6) which contacts the C-terminal tail as well as other internal residues, indicating large-scale changes in the conformation and dynamics of Sox2 in complex with specific DNA.
